# Computational Enhanced Hi-C data reveals the function of structural geometry in genomic regulation

**DOI:** 10.1101/2022.07.12.499232

**Authors:** Yueying He, Yue Xue, Jingyao Wang, Yupeng Huang, Lu Liu, Yanyi Huang, Yi Qin Gao

## Abstract

High-throughput chromosome conformation capture (Hi-C) technique profiles the genomic structure in a genome-wide fashion. The reproducibility and consistency of Hi-C data are essential in characterizing dynamics of genomic structures. We developed a diffusion-based method, C_T_G (Hi-C To Geometry), to deal with the technical bias induced by insufficient sampling in sequencing and obtain reliable gemeotric information of the chromatin. C_T_G properly quantifies dubiously weak or even undetected interactions and produces a consistent and reproducible framework for the 3D genomic structure. C_T_G allows for a reliable genome-wide insight on the alteration of genomic structures under different cellular conditions and reveals correlations between genomic-proximal genes at both transcriptional and translational levels. Cell-specific correspondence between gene-gene and corresponding protein-protein physical interactions, as well as that with the transcription correlation reveals the coordinated inter-molecular structural and regulatory information passage in the central dogma.

## Introduction

The three-dimensional architecture of chromatin is crucial to the functionality of one-dimensional DNA sequences (Oudelaar and Higgs, 2021). However, the concrete correlation between the 3D architecture and its function in genome regulation has not been completely resolved. High-throughput chromosome conformation capture (Hi-C) technique (Lieberman-Aiden et al., 2009) allows for genome-wide profiling of chromatin interactions in 3D-space by performing unbiased DNA-DNA proximity ligation. Hi-C reveals a hierarchical organization of chromatin (Rowley and Corces, 2018) and the 3D architecture is demonstrated to be involved in critical biological processes, such as gene regulation, cell fate decisions, and even evolution (Bonev and Cavalli, 2016). Sharing fixed genetic inheritance, the primary domains that make up the hierarchical organization, such as compartments and topologically associating domains (TADs) are largely conserved across cell types (Rao et al., 2014). On the other hand, the variations of chromatin structures among different cell states are pertinent to their distinct genomic function (Bonev and Cavalli, 2016). Various types of genomic changes are relevant to genetic disorders and can lead to genomic diseases such as cancer (Corces and Corces, 2016; Li et al., 2020). Hence, it’s essential to study the dynamics of chromatin structures, quantifying the variations with cellular states and understanding their functions.

The great success of Next Generation Sequencing (NGS) technology makes it possible to obtain Hi-C data with high throughput. However, the quality and reproducibility of raw Hi-C data are affected by technical and biological bias, and the characterization of the genomic geometry requires normalization tools. A number of normalization algorithms have been developed to remove unwanted systematic bias. The normalization algorithms fall into two main categories: explicit-factor correction and implicit matrix balancing. Explicit-factor correction algorithms such as Hi-C-Norm (Hu et al., 2012) propose parametric models to depict known bias such as GC content, fragments length, and mappability. Implicit matrix balancing algorithms, such as iterative correction and eigenvector decomposition (ICE) (Imakaev, Maxim; Fundenberg, Geoffrey; Patton McCord, Rachel; Naumova, Natalia; Goloborodko, Anton; Lajoie, Bryan R.; Dekker, Job; Mirny, 2012), Knight and Ruiz’s algorithms (Hu et al., 2012), and chromoR (Hu et al., 2012), assume equal visibility for all genomic loci and balance row and column sums. These methods remove reoccurring biological bias and improve the reproducibility of replicated datasets, but leaving unpredictable technical biases unaddressed. The unpredictable technical bias mainly comes from insufficient sampling, resulting in dubiously weak contact strengths and random noise. The correlation between raw matrices and matrices normalized by different algorithms increases with the sequencing depth (Han and Wei, 2017), indicating the importance of sufficient sampling. The randomly directed noise conceals the real biological proximity information and impedes the characterization for variations of the chromatin structures among different cell states. There are multiple computational methods (Djekidel et al., 2018; Lun and Smyth, 2015; Stansfield et al., 2019) aimed for making statistically-grounded comparisons between Hi-C datasets and quantifying statistically significant dynamic changes. A few of them, including diffHiC (Lun and Smyth, 2015) and multiHiCcompare (Stansfield et al., 2019), conduct across-sample normalizations to improve their performances to quantify consistent differential chromatin interactions. The across-normalization methods reduce the random noise, but the problem on intrinsic insufficiency in sampling is not addressed, limiting the performances of these statistically-grounded methods.

The distance matrix is naturally a full matrix and a corresponding contact matrix can be recovered from the distance matrix following a power law approximation, where the strengths of weak or even undetected interactions are properly quantified. Here, we propose C_T_G (Hi-C To Geometry), a diffusion-based algorithm, to treat the technical insufficiency and uncover the geometric structure from Hi-C data (Figure 1). C_T_G takes Hi-C contact matrix normalized by ICE as the input, and outputs a C_T_G distance matrix. The main inspiration of C_T_G algorithm stems from the physical succession of the genomic structure. In perspective of a proximity network, the proximal genomic regions should share similar diffusion manners. The C_T_G distance between pairwise genomic regions is quantified by their genomic-wide diffusion manners and therefore reduce the impact of insufficient sampling for any individual interaction. C_T_G, as a distance-like measurement, allows for genome-wide insight into the correlations between proximal genes in genomic structure and we investigated the correspondence at transcriptional and translational levels.

**Figure 1.**
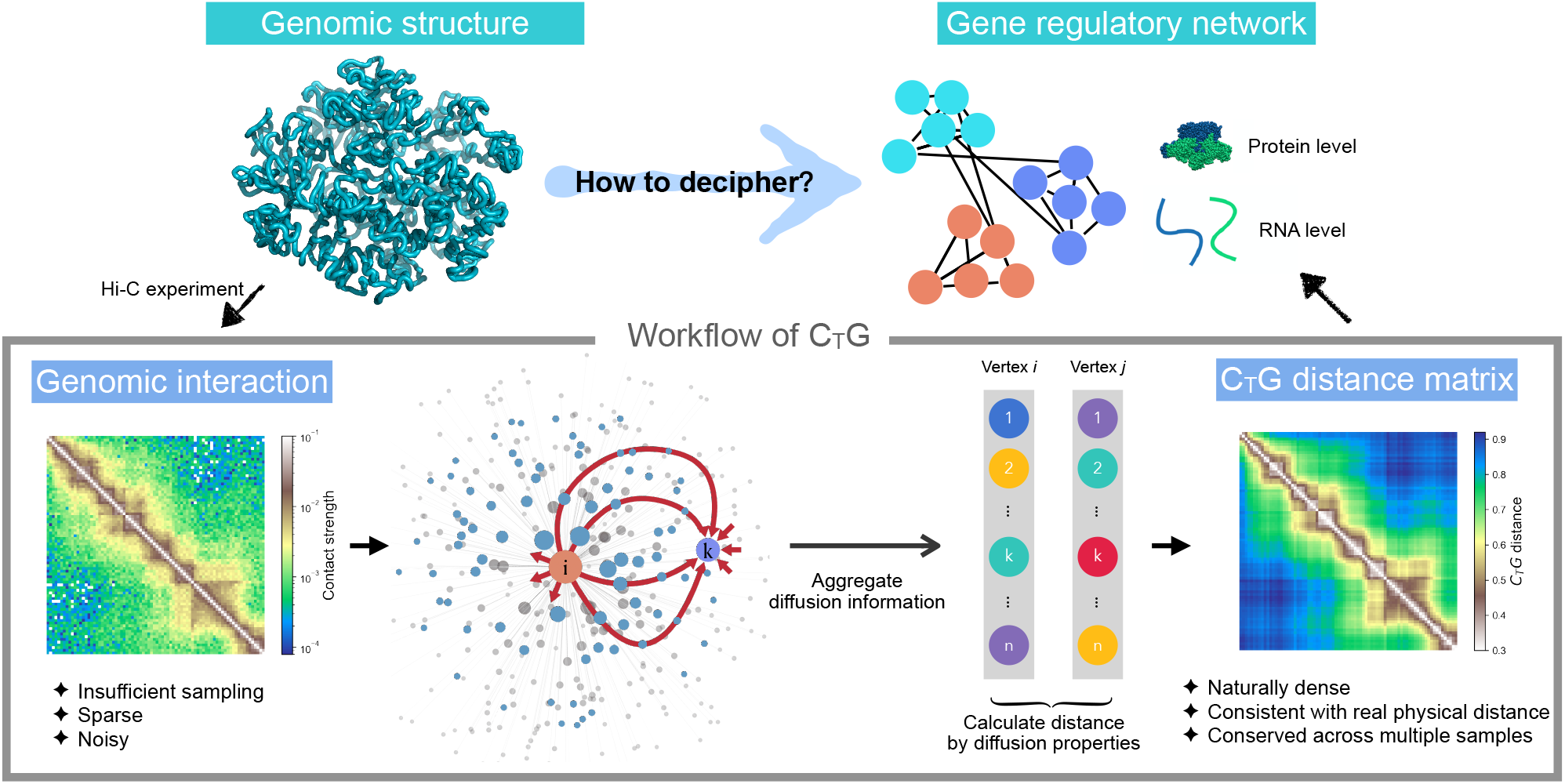
Schematic overview of C_T_G. C_T_G uses a diffusion-based strategy to uncover the gemetry of genomic structure from Hi-C data. C_T_G quantifies the diffusion property of each vertex by aggregating global diffusion information from the vertex to other verteces respectively. And the C_T_G distance between pairwise verteces is calculated by similary of their diffusion properties. CTG allows for a genome-wide insight deciphering the gene regulation information coded in genomic structure.

## Results

### Overall design of C_T_G

The Hi-C contact map depict a proximity network *G*(*V,E*), where the vertices *V*={*v*_*1*_,*v*_*2*_,*…v*_*n*_} denote the non-overlapping genomic regions and the edges *E* = {*e*_*i,j*_} denote the contact strength between pairwise connected genomic regions. Similar to diffusion-based methods for network denoising (Cao et al., 2013; Wang et al., 2018), a Markov prosses (2007) is used to describe the diffusion process on this network. 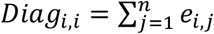, is the element of the diagonal degree matrix *Diag* for the network. The vector 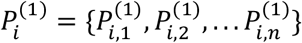 is the conditional transition probability transiting from vertex *v*_*i*_ to *V*={*v*_*1*_,*v*_*2*_,*…v*_*n*_} in one single step. Likewise, 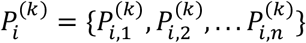 is the conditional transition probability in *k* steps and 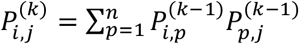. With increasing *k*, the transition probability from *v*_*i*_ to *v*_*j*_ gradually integrates neighbor information and expand the inclusion of edges, since *v*_*i*_ and *v*_*j*_ may not be connected in one step but they can be connected in some finite steps as the network *G* is a connected graph. Taking *k*=2 and 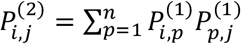 as an example, when the two pairs of vertices (*v*_*i*_ and *v*_*p*_, *v*_*j*_ and *v*_*p*_) are pairwise neighbors, which means 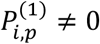 and 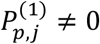, *v*_*p*_ contributes to 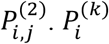 converges to an invariant distribution for connected graph and the difference between 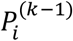 and 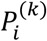 decreases.

It is thus appropriate to use the integrated information on 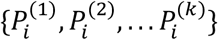 to describe the diffusion manner of vertex *v*_*i*_ within some given number of *k* steps, which can be infinite. In practice, we found that 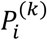 converges rapidly and therefore used the exponential decay to fit the convergence. 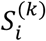 is defined as the weighted summation of 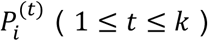:

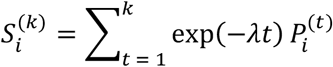

When *k* reaches infinity, 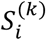 converges to *S*_*i*_ (Supplementary note). As the weighted summation of 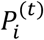, *S*_*i*_ naturally integrates neighbor information of the connected graph, and therefore alleviates in a physics-based manner the problems caused by the Hi-C data sparsity. On the other hand, the exponential decay ensures that the integration does not eliminate the distinction of each vertex, taking the rapid convergence of 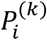 into consideration.

The physical succession of the genomic structure suggests that the proximal genomic regions should share similar diffusion manners. The similarity between pairwise vertices *v*_*i*_ and *v*_*j*_ is quantified by L1 distance between *S*_*i*_ and *S*_*j*_. L1 distance is used as a measure since it mitigates the impact of outliers caused by distance metrices of higher-order terms. A C_T_G distance matrix is then constructed based on the Hi-C contact map. We demonstrate below that C_T_G distance is relevant to real spatial distance and thus provides information on the geometry of the genome. Meanwhile, to fit the contact probability, a C_T_G contact matrix is converted from the C_T_G distance matrix by making use of a power law, according to the power-law dependencies derived from polymer-like behavior (Halverson et al., 2014; Lieberman-Aiden et al., 2009). With the power of 4, the distribution of the reconstructed contact frequency is most similar with raw HiC contact datasets.

### Validation of C_T_G

One way to test whether the sequencing-based method such as Hi-C can faithfully reproduce geometric structure information is to make comparison with fluorescence in situ hybridization (FISH) imaging data (Su et al., 2020), as the latter provides direct spatial position information of individual loci. Ref. Su et al., 2020 provided high-resolution imaging data on the coordinates at 50-kb resolution for Chr2 and Chr21 of human lung fibroblast (IMR-90) cells. The median spatial distance between pairs of imaged loci is thus a physical distance measurement (Figure 2A and 2B, right pannel). Taken the Hi-C data of IMR-90 (Rao et al., 2014), one can perform a direct comparison between the spatial distance and the inverse contact probability and the Pearson correlation coefficient is 0.790 and 0.897 (with logarithm transformation) for Chr2 and Chr21, respectively, which is to some extent satisfactory. In contrast, as shown in Figure 2C, the calculation of C_T_G distance matrix (Figure 2A and 2B, left pannel) improves its linear correlation with the physical distance measurement and the corresponding Pearson correlation coefficient with the median spatial distance matrix reaches 0.952 and 0.930, respectively. These results show that the C_T_G method provides a more accurate calibration between two different experimental methods and the distance metrics generated by the C_T_G method reproduces that observed by super-resolution experiment.

**Figure 2.**
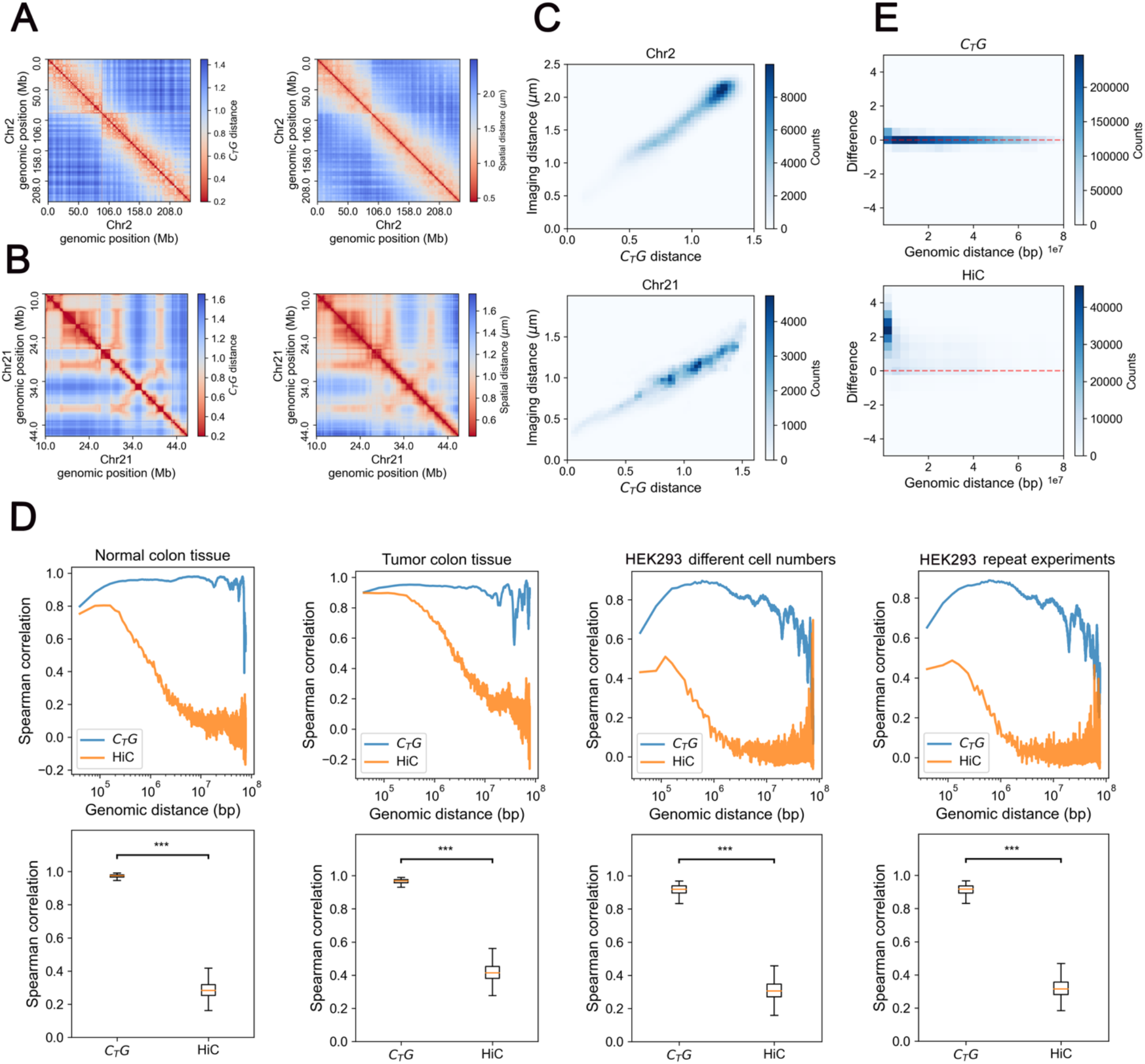
Validation of C_T_G. (**A**) The C_T_G distance matrix (left) and the median spatial distance matrix (right) of chr2 (resolution of 50kb). (**B**) The C_T_G distance matrix (left) and the median spatial distance matrix (right) of chr21 (resolution of 50kb). (**C**) the correlation between CTG distance matrix and the median spatial distance matrix of chr2 and chr21. (**D**) The Spearman correlation for genomic sequence distance (upper pannel) and for individual genomic region(lower pannel) between pairwise contact matrices derived from 1) normal colon tissue samples; 2) tumor colon tissue samples; 3) different numbers of 293 cells; 4) repeated experiments on 293 cells. *** represents P-value<10^−300^ (t-test). (**E**) The MD plots between two normal colon tissue samples in view of genomic sequence distance.

Next, we evaluate the robustness of C_T_G contact propensity map by applications to different samples and compare the Hi-C data derived from a) normal colon tissue samples of different individuals (Johnstone et al., 2020), b) tumor colon tissue samples (Johnstone et al., 2020), c) different numbers of HEK293 cells (sample 0923-2 and 0923-4), d) repeated experiments on HEK293 cells (sample 0923-4 and 1002-5). The robustness of C_T_G is assessed by calculating Spearman correlation coefficient of spatial interactions from different samples at various genomic distances. Such a calculation is equivalent to calculating Spearman correlation coefficient of diagonal elements of Hi-C maps. For an Hi-C contact map treated after ICE normalization, the correlations between different samples decrease sharply as genomic distance increases (Figure 2D, upper pannel), indicating that the normalized Hi-C contact map is of high confidence level at scales up to about 5Mb but not longer. In contrast, the correlations of C_T_G contact maps are significantly higher and hardly decrease with the genomic distance. We also compared the Spearman correlation coefficient for individual genomic regions between Hi-C and C_T_G contact maps, equivalent to calculating Spearman correlation coefficient of each row of different contact maps (Figure 2D, lower pannel), where the latter also display a higher consistency than the former. In addition, the systematic bias between different datasets for Hi-C and C_T_G contact map were quantified by a MD plot (Minus, or difference vs. Distance plot) (Stansfield et al., 2018), to visualize the differences between two datasets accounting for the linear genomic distance between interacting genomic regions. *M* is defined as the fold-change between two Hi-C datasets, with its element *M*_*ij*_ *= log*_*2*_*(IF*^*1*^_*ij-*_ *IF*^*2*^_*ij*_), where *IF*^*1*^_*ij*_ and *IF*^*2*^_*ij*_ are contact strengths between pairs of genomic regions from two datasets. *D* is defined as 1D genomic distance of pairwise genomic regions. In this was, the systematic bias between different datasets is reflected by the deviation of *M* from the *M*=0 baseline. The MD plot (Figure 2E) of C_T_G contact map is approximately symmetric about *M*=0 baseline without any prior fitting. In contrast, for the Hi-C contact map, only 30% non-zero elements can be faithfully calculated due to the limitation of sparse data. The distribution obtained for the Hi-C contact map (Figure 2E, lower panel) deviates significantly from the baseline, indicating the impact of systematic bias.

We note here that the unprocessed Hi-C contact map is subject to large noise due to incomplete statistics, and the large variance of long-range interactions (>5Mb) among similar samples indicates that weak interactions or long-range interactions tend to be unreliable. Therefore, a genome-wide comparison between different Hi-C datasets is ambiguous, due to the noisy and sparse data. By incorporating the genome-wide diffusion property of each genomic regions into consideration, the problem associated with insufficient sampling for singular interactions is sufficiently corrected. The C_T_G contact/distance maps reveal the hidden reproducibility of Hi-C data and more importantly, that the putative topologies of genomic structures are conserved across different cell numbers and even different individuals. The genomic structures recovered by C_T_G algorithm thus allow for direct comparison for replicate experiments and even for samples from different individuals/experimental setups. Such a property of C_T_G makes it suitable for characterizing the changes of genomic structures under different conditions.

### C_T_G characterizes the global structural changes in Colorectal Cancer pathogenesis

In this section, we use the C_T_G method to analyze genomic structures derived from normal and tumor colon Hi-C data. Compartmental recognition was performed in a previous study (Johnstone et al., 2020) on these datasets, which associated the compartment changes during colorectal cancer pathogenesis with stemness, invasion, and metastasis of tumor. In the following, we show that C_T_G allows for new insights into cancer-related changes of genomic structure. To ensure the consistency and reproducibility of our analysis, pairwise normal and tumor samples derived from 4 individuals were compared. We took chromosome 17 as an example in our latter single chromosome analysis to simplify our discussion. The conclusions are the same for other chromosomes.

As can be seen from Figure 3A, the overall pattern of C_T_G distance matrices clearly distinguishes normal from tumor colon samples. From direct visualization, the fine plaid patterns of normal samples become significantly blurred in cancer, where the distinct genomic “chess-like squares” are no longer properly segregated and the specific long-range aggregation weakens. To be more quantitative, we calculated the contrast ratio of the genomic “squares” over their proximal neighbors (Figure 3B, Method). The contrast ratios were found to be significantly higher for normal samples than tumor samples (P-value=0.0084) and were conserved across 4 individuals. Such a result indicates that there is a clear insulation between neighboring regions in normal tissues, the strength of which weakens in cancer samples. This change in genome insulation indicates the potential transcriptional dysregulation in carcinogenesis. One important factor affecting genome insulation is CTCF. It is known that CTCF/cohesin-binding sites are frequently mutated in cancer (Katainen et al., 2015) and immortalized cancer cell lines display a low CTCF occupancy with the hypermethylation of CTCF/cohesin-binding sites(Ong and Corces, 2014). However, it was also reported that the compartmentalization of mammalian chromosomes were independent from CTCF (Nora et al., 2017). The observations on mutli-scale chromatin structure changes thus suggest the influence from systematic aberration such as the uncontrolled cell cycle (Hanahan and Weinberg, 2000) in addition to the absence of chromosomal structure regulator, such as CTCF. Such a possibility has been suggested by Ma et al., 2015.

**Figure 3.**
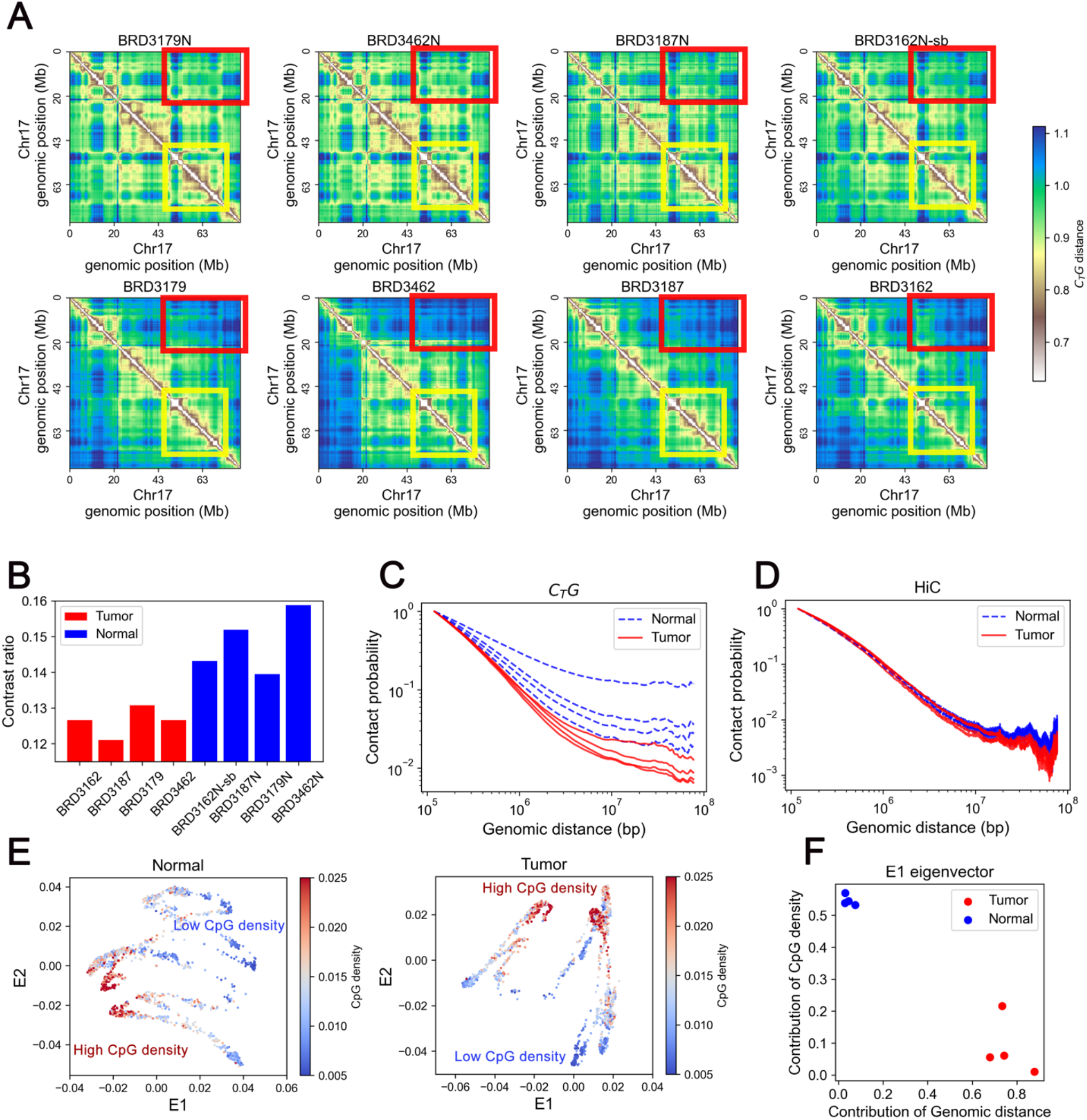
Global structural patterns of Colorectal Cancer revealed by C_T_G. (**A**) The C_T_G contact maps for normal (upper pannel) and tumor (lower pannel) colon samples. Each column represents pairwise normal and tumor samples derived from the same patient. The yellow and red squares are examples of the differences between normal and tumor samples. (**B**) The contrast ratio of the C_T_G distance map, the blue bars correspond to normal samples and the red bars correspond to tumor samples. (**C**) Contact probability as a function of genomic distance calculated from the C_T_G contact map. (**D**) Contact probability as a function of genomic distance calculated from the Hi-C contact map. (**E**) The 2D Laplacian Eigenmaps of C_T_G distance matrices for pairwise colon normal and tumor samples. Each point represents a 40kb genomic region. The color is used to represent the CpG density of the corresponding genomic region. (**F**) Contribution of sequences properties to structure-related E1 eigenvector.

Next, we calculated the reconstructed contact as a function of the 1D genomic distance (Figure 3C). It can be seen that the tumor samples display large decay rates in ∼Mb region and the comparison between normal and cancerous C_T_G distance matrices suggests the loss of specific long-range interactions in colon cancer, as revealed by Figure 3C. In comparison, the decay curve derived from Hi-C data normalized by ICE only varies more significantly over different sample paris (Figure 3D), again validating the effectiveness of C_T_G in revealing the consistent difference between normal and cancer cells.

Sequence properties, especially CpG density, was reported to be an important factor affecting the organization of genomic structure (Liu et al., 2018). To gain understanding on how one-dimensional DNA sequences affect the organization of three-dimensional genomic structure, we performed dimensionality reduction on C_T_G distance matrix. The non-linear Laplacian Eigenmaps (see Methods) was employed for dimensionality reduction, as the eigenvectors obtained by this method are interpretable and reveals information on hierarchical clustering (Figure 3E, Figure S1). Sorted by eigenvalues, the leading eigenvector E1 reflects the predominant structural patterns. We quantified the contribution of sequence properties, including sequential similarity (CpG density) and sequential distance, to genomic structure, by projecting the structure-related eigenvectors on these sequence properties. Reflected by projection of E1(Figure 3F), the dominant factor in structure determination changes from sequential similarity in normal cells to sequential distance in colon cancer, affecting the organization of A and B compartmental domains and probably resulting in the dysregulation of transcriptionally active or inactive states (see Discussion).

### CTG reveals the coupling of co-expression and genomic proximity during Colorectal Cancer pathogenesis

The genomic structure is believed to play a crucial role in the precise gene expression program (Elimelech and Birnbaum, 2020; Oudelaar and Higgs, 2021). The genomic interactions between gene promoters and distal cis-regulatory elements have been studied extensively (Li et al., 2022). Since less attention has been paid on the function of gene-gene co-localization in genomic structures, we investigate here the physical gene-gene interactions at genomic levels, represented through the contact between genomic bins in 40 Kb resolution which contain these genes. Of special interest is whether a correlation exists between gene-gene contact in chromatin and gene co-expressions at the transcript level. The correlation network at transcript levels was characterized by spearman correlation coefficients of RNA-seq data, with the RNA-seq data derived from the Cancer Genome Atlas (TCGA) program, for 86 pairwise normal and tumor colon samples. The interaction network at genomic levels was quantified by C_T_G distances. The two networks were aligned together in perspective of the genomic position of each gene.

The overall patterns of co-expression matrix changed significantly from normal to tumor samples (Figure 4A and 4B). Taking chromosome 17 as an example, the expression correlation between pairs of genes detected in tumor samples decreases sharply as a function of the corresponding linear genomic distance between the gene pairs increases, whereas this function barely changes with the genomic distance in the case of normal tissues (Figure 4C). Intriguringly, the dependence of co-expression on the genomic distance resembles that of C_T_G contact map, implying a potential relationship between genomic proximity and co-expression. To be more specific, we evaluated the one-to-one correspondence between genomic co-localization and co-expression. For both tumor and normal samples, the proximal gene-pairs tend to co-express at the transcript level (Figure 4D), and such an inter-dependence is stronger for tumor than normal samples. In reverse, gene-pairs that share a similar expression pattern tend to be proximal at genomic levels for tumor samples (Figure 4E), which is again more prominent for tumor than for normal samples. Such a difference between tumor and normal samples indicates an increased correlation between genomic structure and gene transcription in cancers in perspective of gene-gene interplay. Compared to cancer sample, there is a weaker correlation between gene pair proximity and their expression correlation across normal samples, for which genes of large linear and spatial distances can be highly correlated in expression, suggesting more important roles of regulation mechanisms besides spatial co-transcription, such as histone modification or DNA methylation, in normal cells than in their cancerous counterparts. The elevated dependence of gene co-expression on their spatial interaction in chromatin may suggest that the gene expression regulation becomes more directly correlated with genomic structure. Interestingly, it was discovered recently that the RNA and protein levels become more strongly correlated in carcinogenesis, supporting that the regulation network simplifies in cancer pathogenesis (Nusinow et al., 2020). Moreover, besides solid tumors, we also found similar correspondence of gene-gene proximity and gene co-expression in acute lymphoblastic leukemia samples (Figure S2).

**Figure 4.**
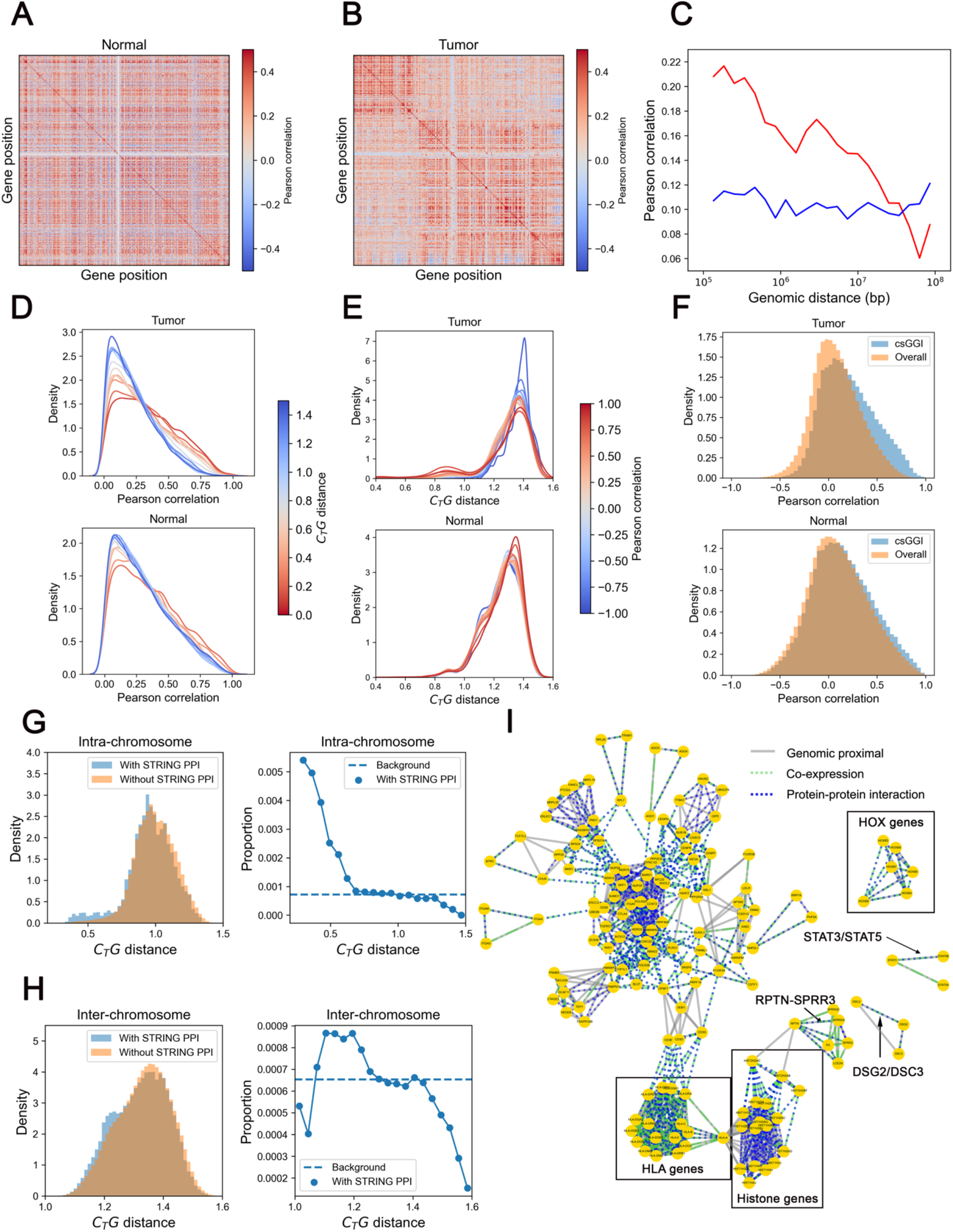
Passage of gene-gene interplay from genomic level to transcription and protein levels in Colorectal Cancer. (**A**) Gene-gene transcriptional Pearson correlation matrix of chromosome 17 of normal colon samples. (**B**) Gene-gene transcriptional Pearson correlation matrix of chromosome 17 of tumor colon samples. (**C**) The averaged correlation coefficients as a function of 1D genomic distance between gene pairs. (**D**) The distribution of transcriptional Pearson correlation under different C_T_G distance of whole chromosome, the color of each line indicates corresponding C_T_G distance. (**E**) The distribution of C_T_G distance under different Pearson correlation of whole chromosome, the color of each line indictes corresponding Pearson correlation coefficient. (**F**) The distribution of correlation of gene-pairs with csGGIs and overall background (**G**) The distribution of C_T_G distance between intra-chromosomal gene pairs with and without STRING PPIs for whole chromosome in tumor sample (left pannel); The proportion of intra-chromosomal gene-pairs with STRING PPI at different C_T_G distances in tumor sample (right pannel). (**H**) The distribution of C_T_G distance between inter-chromosomal gene pairs with and without STRING PPIs for whole chromosome in tumor sample (left pannel); The proportion of inter-chromosomal gene-pairs with STRING PPI at different C_T_G distances in tumor sample (right pannel). (I) The gene network integrating colon cancer-related gene-gene interplay at DNA, RNA and protein levels. The three kinds of edges indicate gene-gene interplays at three levels.

Next, we analyzed the local spatial contacts in chromatin for individual genes (see Methods), where spatial gene-gene interactions (GGIs) are characterized. The interactions formed in cancer but not in normal tissue are referred as cancer specific GGIs (csGGIs). Noticeably, genes involved in csGGIs are prone to be more positively correlated in tumor samples than normal samples comparing with respective background (Figure 4F). These csGGIs tend to be properly insulated in normal cells but not in cancer. We expect the csGGIs in genomic structures of tumor colon samples quantified by C_T_G algorithm to play an important role in transcriptional co-regulation between genes. Therefore, we further select csGGIs with notable changes in RNA correlation (tumor correlation >0.5 and normal correlation <0.1, the criterion is robust) and construct a csGGIs network. We found that the cancer-related genes (see Methods) are indeed enriched in the network, as 4.33% genes involved in this network are cancer genes with 0.28% of all coding genes being cancer genes. The cancer genes, including ERBB3, HRAS, MAP2K2, PTK6, RAC1, SDC4, TSC2, SRC, among others, are connected with more than 5 genes and thus may play central roles in this network (Figure S3-S5). Meanwhile, most of them are reported to be highly relative in colorectal cancer pathogenesis (Liu et al., 2021; Serebriiskii et al., 2019; Wang et al., 2021). Deciphering the gene-gene interaction and resulted changes in regulation networks is expected to render further understanding on the specific functionality of these genes in addition to that provided merely by mutation of single genes. As the cancer genes are inferred from only cancer-related mutations, we next performed functional annotation analysis on all genes connected with more than 5 genes in this network (Table S1) and found these genes to be strongly involved in epidermal growth factor receptor (ERGF) signaling pathway and proteoglycans in cancer. In addition, HRAS, RAC1, SOS2, MAPK3 and MAP2K2 directly participate in colorectal cancer KEGG pathway. HRAS is involved in multiple cancer-related process and genes interacting with HRAS in cancer genomic structure, for example, IFITM3, DRD4, IRF7 and NLRP6, are heavily involved in immune response. Such an analysis likely provides a new perspective on the roles of immune responses in cancer pathogenesis.

### C_T_G reveals the information passage from genomic proximity to protein-protein interaction in colorectal cancer pathogenesis

After interrogation on the interplays between gene-pairs at DNA levels and their transcript product, we ask whether such information is further passed along the central dogma, such that gene-gene interaction at the chromatin level affects the interaction between their translational products. The interplays at protein levels were evaluated by physical protein-protein interactions derived from the STRING project (Szklarczyk et al., 2021). The genomic interactions and PPIs were aligned by genes and protein isoforms generated from corresponding genes. As shown below, we did identify associations between genomic structure and protein-protein interactions (PPIs) in both normal and tumor samples that have not been discussed before.

First, it can be seen from left panel of Figure 4G, 4H, S6A and S6B that the C_T_G distances between gene-pairs with their proteins forming known/predicted PPIs tend to be more proximal than those without PPIs, for both intra-chromosomal gene-pairs with a more stable genomic structure and inter-chromosomal gene-pairs with a more flexible genomic structure. We also calculated the proportion of gene-pairs containing PPIs under varied C_T_G distances and show the results in right panel of Figure 4G, 4H, S6A and S6B, from which one observes that the spatially proximal gene-pairs are more likely to have their product proteins to form PPIs. These results suggest that contact information deposited in genomic spatial structures has a tendency to pass to the protein level. Since the information passage of DNA-DNA (gene-gene) interaction to protein-protein interaction inevitably goes through RNA, we next examined the correlation between different genes at RNA and protein levels. Interestingly, gene pairs forming PPIs in the STRING dataset are indeed more prone to be correlated in transcription than randomly chosen pairs and such a tendency is found across different tumor types (Figure S7). Although co-expressions are a portion of gene interplays at RNA levels and PPIs in the datset are not tissue-matched, gene pairs with GGIs and PPIs are more correlated in transcription than those only with PPIs (Figure S8). Such results suggest that the information of gene regulatory network is at least partially coded in 3D genomic structures and tranferred to RNA and protein levels along with the central dogma, in a way beyond correct coding and functioning of single genes, but also at the message-passage level in form of gene-gene interations.

Integrating gene-gene interplay at DNA, RNA and protein levels, a number of gene pairs are seen to be at the center of interaction network for colon cancer (Figure 4I). For example, STAT3/STAT5, DSG2/DSC3, and RPTN/SPRR3, all possess genomic proximity, transcription coregulation and potential protein interactions inferred from STRING. In fact, these genes are all reported to be involved in colorectal tumorigenesis. For example, STAT3 are known biomarkers for colon cancer as it is necessary for proliferation and survival in colon cancer-initiating cells (Lin et al., 2011), and STAT5 are reported to be involved in regulation of colorectal cancer cell apoptosis (Du et al., 2012). The downregulation of DSG2 and DSC3 in colon cancer cells was found to suppress colon cancer cell proliferation (Cui et al., 2011; Kamekura et al., 2014), and DSC3 is involved in tumor suppression activity (Cui et al., 2019). Finally, the overexpression of SPRR3 is known to promotes cell proliferation through AKT activation (Cui et al., 2011). The interactions between multiple genes can also be observed in the chromatin structure. For example, close proximity is seen among HLA (Human Leukocyte Antigen) genes (Figure S9). It is known that the relavent translational products make up the HLA class I (HLA-A, HLA-B, HLA-C) and class II (HLA-DQ, HLA-DR) complexes, which play important and distinctive roles in presenting processed peptide antigene (Choo, 2007; Giudizi et al., 1987). The results indicated that not only direct protein interations within each class of complex, but also co-regualtion between the two complexes may be partially coded in genomic structure, although they are distant in the linear genome.

### C_T_G reveals the tissue-specific coupling of protein-protein interaction and genomic interactions

The integrated STRING PPI dataset contains both tissue-matched and unmatched PPIs, which allows the statistical anaylsis on GGI-PPI correlation but limits one from precisely match GGIs with PPIs in a cell-state specific manner. To overcome this limit, we next performed analysis based on the tissue-matched PPI datasets from the Affinity-Purification Mass Spectrometry (APMS) technique (see Methods). Fortunately, BioPlex project has compiled a comprehensive dataset of protein-protein interactions of HCT116 cells (Huttlin et al., 2021), which allows us to quantify the correlation between genomic interactions and protein interactions for this colorectal carcinoma cell line. The cell-matched BioPlex PPIs consist ∼71,000 interactions and they are all included in our analysis. Consistent with the results obtained using STRING datasets, as shown in Figure 5A and 5B, genomic proximal gene pairs in HCT116 cell are also more likely to possess corresponding PPIs and on the other hand, gene pairs with corresponding PPIs also tend to be spatially closer in genomic structure than those without known PPIs, although the current PPI list is proabably far from being complete.

**Figure 5.**
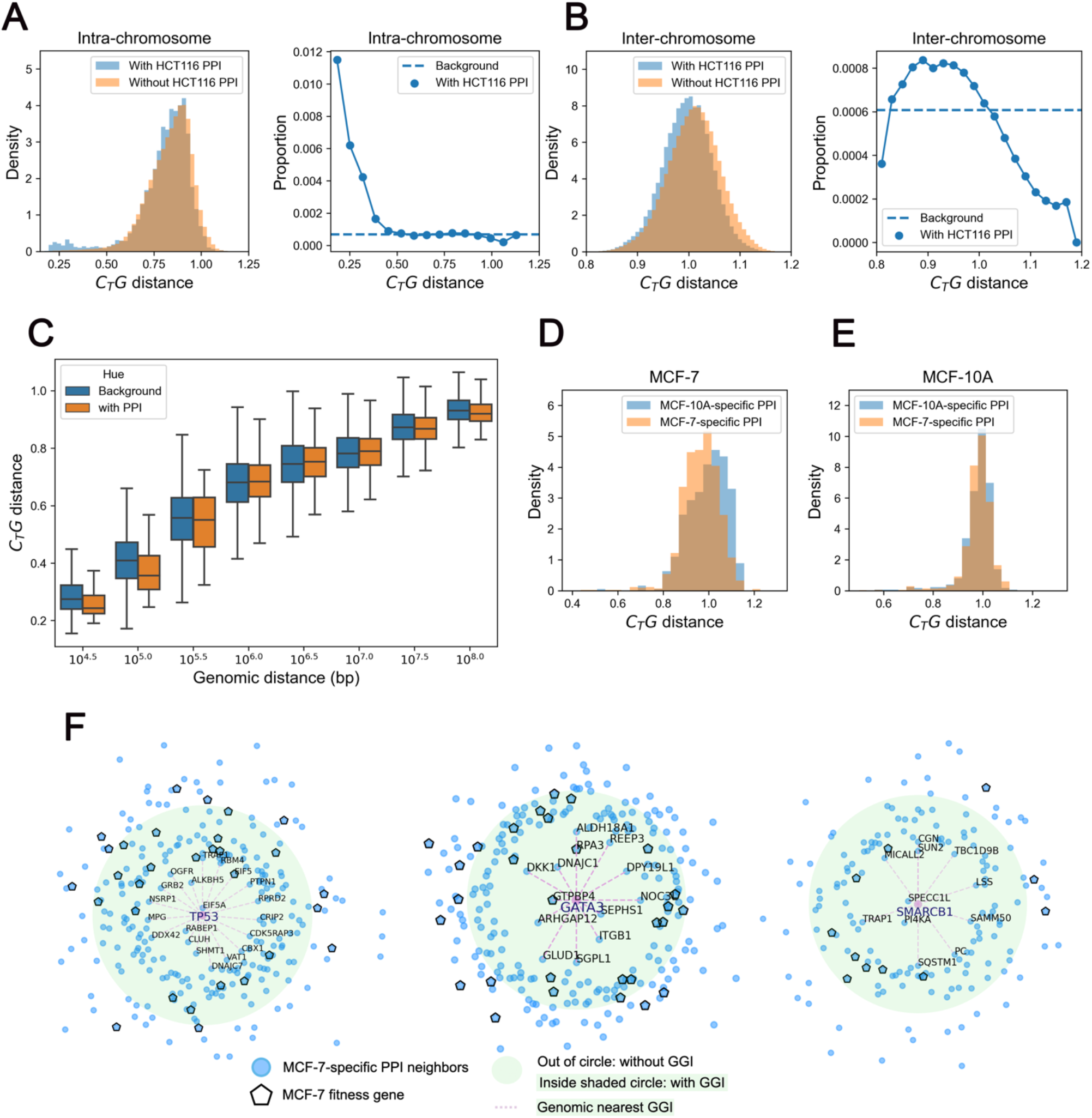
The tissue-specific correspondance of protein-protein interaction and genomic proximity. (**A**) The distribution of C_T_G distance of intra-chromosomal gene pairs with and without HCT116-related PPIs for whole chromosome (left pannel); the proportion of intra-chromosomal gene-pairs with HCT116-related PPI at different C_T_G distances (right pannel). (**B**) The distribution of C_T_G distance of inter-chromosomal gene pairs with and without HCT116-related PPIs for whole chromosome (left pannel); the proportion of inter-chromosomal gene-pairs with HCT116-related PPI at different C_T_G distances (right pannel). (**C**) The C_T_G distance of gene pairs in fixed 1D genomic distance. (**D**) C_T_G distance of gene pairs with MCF-7-specific and MCF-10A-specific PPIs in MCF-7 cell. (**E**) C_T_G distance of gene pairs with MCF-7-specific and MCF-10A-specific PPIs in MCF-10A cell. **(F)** TP53,GATA3, SMARCB1 related MCF-7-specific PPIs,the distance to TP53 indicates the CTG distance and the green circle indicates the background distance,the pink scatter indicates MCF-7 fitness genes and the dashes inducate genomic proximal neighbors.

The mutual correspondence between GGI and PPI uncovers a significant correlation between genomic interactions and protein-protein interactions. The genomic proximity information appears to be partially preserved in both transcription and translation. Furthermore, the intra-chromosomal gene-pairs with PPIs (Figure 5A and 5B, left pannel) displayed a tighter correlation with genomic structure than inter-chromosomal ones (Figure 5A and 5B, right pannel). Interestingly, it is known that genes with related functions tend to cluster along the linear genome and in individual chromosomes (Hurst et al., 2004). The higher intra-than inter-chromosomal DNA, RNA and protein coupling is consistent with this functional requirement. Next, to exclude the impact of 1D genomic distance within chromosomes, we evaluated GGI-PPI correlation at fixed genomic distances and found that gene-pairs with corresponding PPIs tend to be more proximal in all genomic distances (Figure 5C) than those without. Limited by a majority of weak or even undetected interactions, these signals are insignificant in raw Hi-C datasets with 90% zero-elements, again demonstrating the importance of further data processing for Hi-C matrix. We also performed functional annotation analysis for proximal gene-pairs with tissue-matched PPIs (Table S2 and S3). These genomic-proximal intra-chromosomal PPIs significantly correlate with cell adhesion and immune response, enrichend in “interferon signaling pathway” and “antigene presentation” (HLA genes). In accrodance, interferon gene family is heavily involved in cancer-related pathways, such as those of JAK-STAT and PI3K-Akt signaling (Burke et al., 2014; Horvath, 2004). In the meanwhile, HLA genes play vital roles in cancer immunotherapy (Anderson et al., 2021). The interactions of HLA genes in both genomic and protein levels in colon cancer cell line are consistent with findings on solid colorectal cancer samples. On the other hand, the functions of genomic-proximal inter-chromosomal PPIs are relavent with RNA exosome and proteasome which mediate the degradation of RNA and protein (Makino et al., 2013). The degradation system was shown to play important roles in cancer studies (Manasanch and Orlowski, 2017; Taniue et al., 2022) and the two degradatioin systems may follow common principles (Makino et al., 2013). These results demonstrated the possible roles chromatin and corresponding protein complex structures may play for the establishment of cell identity, as the structural-related PPIs are in correspondence with the cell-specific biological processes.

Next, we studied the specific genomic and protein interactions of breast cancer cell line MCF-7 and its normal counterpart MCF-10A cells (Kim et al., 2021), and compared between them. The specific PPIs were quantified by over-expression affinity purification–mass spectrometry (PPI-score>0.65) (Kim et al., 2021). The number of MCF-10A-specific PPIs is 559 and that of MCF-7-specific PPIs is 1325. From Figure 5D, one observes a clear tendency that gene pairs with MCF-7-specific PPIs are more likely to possess genomic interactions in MCF-7 cells rather than MCF-10A-specific PPIs, while in contrast such a trend is insignificant for MCF-10A cells (Figure 5E). In addition, gene pairs with MCF-7-specific PPIs are more distal (t-value = -16.23, P-value = 1.79 × 10^−57^) and those with MCF-10A-specific PPIs are more proximal (t-value = 7.08, P-value = 1,99 × 10^−12^) in MCF-10A cells than in MCF-7 cells. These results thus reflect a tissue-specific correspondence between GGIs and PPIs. The breast cancer cell line MCF-7 displaying a more significant correspondence than its normal counterpart may reflect that fewer cell-specific PPIs were identified in the normal than the cancer cells. This observation may also indicate the cancer-specific PPIs to be more strongly correlated with the changes in genomic structure, although the inference requires more experimental evidence due to the limited quantity of MCF-10A-specific PPIs. As specific and important examples, we analyzed TP53, GATA3, SMARCB1 and their corresponding MCF-7-specific PPIs neighbors. As shown in Figure 5F, the PPI neighbors of these genes, for example, CBX1/TP53, ITGB1/GATA3 and PI4KA/SMARCB1, tend to be proximal judged by comparison to their mean distances to all genes. Interestingly, their proximal PPI neighbors enrich more MCF-7 fitness genes (Behan et al., 2019), such as EIF5/TP53, GTPBP4/GATA3 and PAM16/SMARCB1, than distal PPI neighbors do in genomic structure, suggesting the importance of genomic structure to cell functionality and survivability.

In summary, C_T_G revealed that a proportion of genomic proximity information is directly reflected at both transcriptional and translational levels. Such an observation suggests that the PPI information is at least partially coded through genomic proximity in the nucleus (see Discussion).

## Discussion and conclusions

We present in this paper a computational method, C_T_G, which was shown to significantly alleviate the insufficient sampling problem of Hi-C datasets. C_T_G takes a HiC contact matrix normalized by ICE as input, and outputs a reconstructed distance/contact matrix, enhancing extremely weak or even undetected interactions in a statistically reliable way. The C_T_G distance matrix is naturally a dense matrix and was shown to be highly consistent with imaging data obtained by FISH technique, thus validating the physical interpretation of the former. We next validated the reproducibility and consistency of C_T_G contact matrix using different cell numbers and even across different individuals and quantified the impact of residual systematic bias. Compared to Hi-C dataset upon normalization, C_T_G generates a reproducible and stable framework to characterize the variation of genomic structures among different samples and experiments. Using this method, we characterized the global changes of genomic structures in colorectal cancer pathogenesis and the changes are consistent across samples taken from different patients. The C_T_G distance matrices can be compared among different samples and permit quantification of the chromatin structure changes, including the loss of specific long-range interactions and dysregulation of transcriptional insulation. These changes are distinguished from sequence-related changes such as gene mutations and structural variations (including deletion, duplications, insertions, inversions and translocations).

Dimensional reduction on C_T_G distance map also reveals the sequence dependence of hierarchical chromatin structure. The organization of A and B compartmental domains is tightly correlated with the 1D sequence similarity, with compartment A of high CGI density compartment B of low CGI density (Liu et al., 2018). In colon cancer, the dominant factor in structure determination appears to change from sequential similarity in normal cells to sequential distance, impeding the long-range interactions of compartmental domains with similar sequence composition. Meanwhile, we investigated the potential correlation between genomic structure and transcriptional co-regulation in colon cancer, and found that the dysregulation in RNA-RNA correlation is at least partially encoded in the genomic structure and can thus be decoded by chromatin structure analysis. We therefore believe that the understanding of the genomic structure can provide a deeper insight into cancer progression and therapy.

In fact, the precise gene expression programed through interactions between gene promoters and distal cis-regulatory elements has been widely investigated. The role of 3D chromatin structure in gene expression regulation has been demonstrated through the importance of loop, TAD formation as well as compartmentalization, although significant uncertainties remain. C_T_G allows for a genome-wide interrogation on the correlations between proximal genes in genomic structure and their functions at transcriptional and translational levels. According to the central dogma, the sequence information of the DNA is mapped into that of RNA and then proteins, effectively resulting in a passage of the one-dimension coding information. From a chemical point of view, the central dogma maps the chemical formula of DNAs to RNAs, and then to proteins, at a single molecule-level and in terms of individual genes. We found here that the flow of information in the central dogma is also manifested as the transmission of gene-gene interplay information, where genomic gene-gene interactions at DNA levels are correlated with co-expressions at RNA levels and protein-protein interactions at protein levels.

Firstly, from DNA to RNA levels, genome-wide correspondence between genomic proximity and co-expression in colorectal cancer was detected in this study. Such an observation triggers us to speculate that the long-range interactions of genomic structure plays a fundamental role in the global transcriptional regulation, ensuring that specific linearly distant genes can share similar transcription environment, such as transcription factor binding and epigenetic hallmarks, and thus are co-regluated. Secondly, from DNA to protein levels, the associations between genomic proximity and PPIs were also detected. We demonstrated such an association on both integrated and tissue-matched PPI datasets. The genomic-proximal PPIs were found to be enriched in tissue-specific biological processes in several cell lines with available data, including HCT116, MCF-7 and MCF-10A. Thirdly, from RNA to protein, it is confirmed that gene pairs with detected PPIs are prone to be positively correlated in transcription for various types of normal and cancer samples deposited in TCGA. Hence, a more comprehensive picture on the biological information passage through central dogma thus likely goes beyond the single gene (protein) and the sequence (chemical formula) level and includes more complex interaction information (Figure 6). In this scenario, the three layers of regulatory networks (roughly speaking, DNA, or more precisely, chromatin, RNA, and protein) are inter-connected not only at the single gene level but also partially at the levels of gene-gene and protein-protein pairs. In this sense, not only genetics but also epigenetics information is passed through DNA to proteins. The distinct epigenetic hallmarks affect the accessibility and TF and RNA binding preference to DNA of specific genomic regions, and introduce distinct gene-gene interactions over similar 1D DNA sequence for different tissues. These interactions are all likely to participate in the establishment of tissue-specific gene regulatory networks.

**Figure 6.**
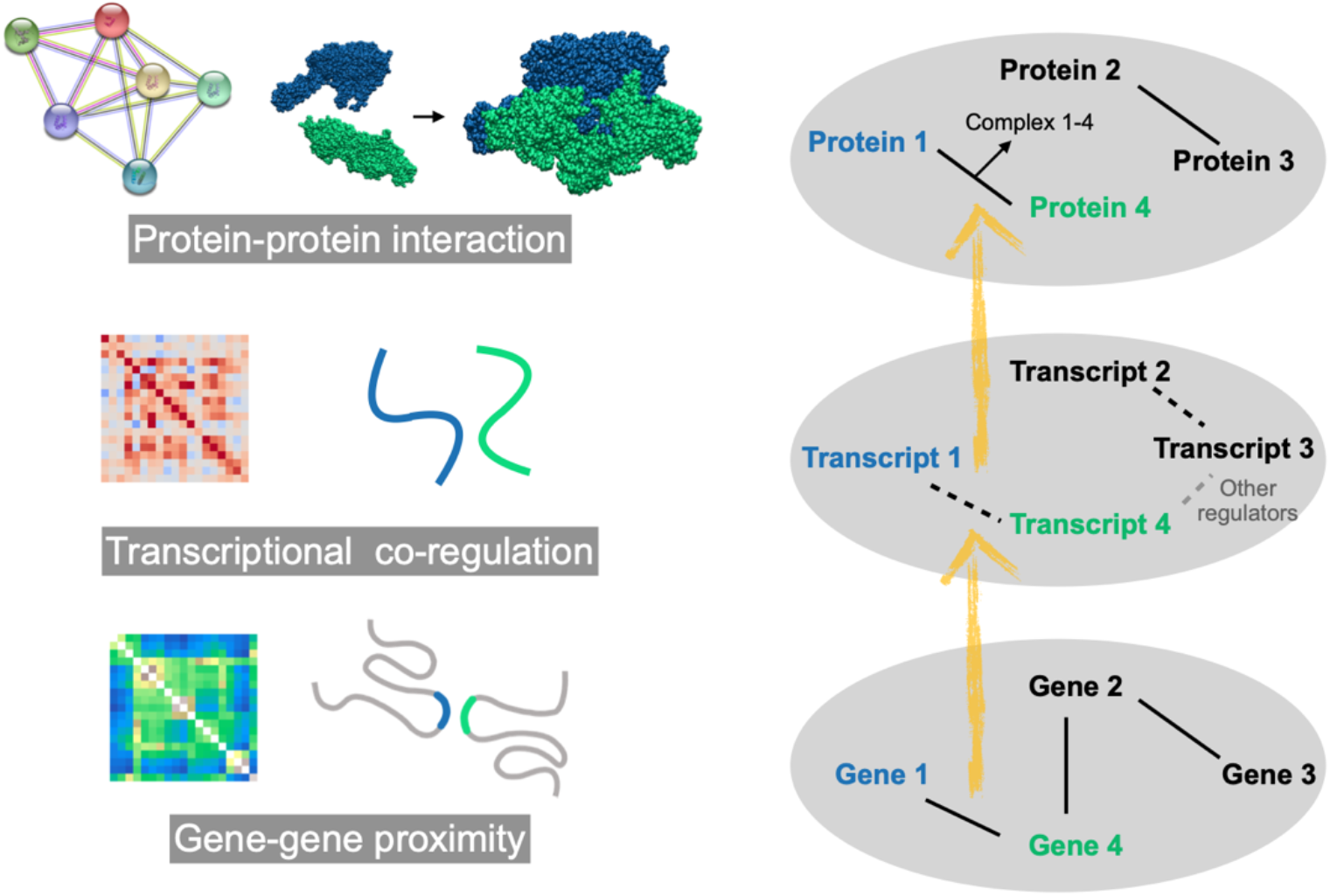
Passage of gene-gene interplay through central dogma. Lines between pairwise genes, transcripts and proteins represent gene-gene interaction, transcriptional co-regulation and protein-protein interaction, respectively.

The storage and passage of interactome information in genomic structure is crucial for tissue-specificity and stability of the regulatory networks. The tissue/cell-specific protein-protein interaction play essential roles in functional orgnization of regulatory networks (Huttlin et al., 2021). However, proteins can vary heavily in number of copies, diffuse relatively freely in the cell if not anchored, and can have short lifetimes. Many of them are also required to respond quickly to external signals and other changes of cellular states. The cell is painfully crowded and complexed for proteins to find and associate with each other faithfully in a timely and well-organized manner, as required by signal transduction, especially if the population and distribution of individual proteins is entirely random or independent of each other. The highly responsive protein-protein interactions also impose difficulties for the proteins to maintain cell state-related information with constant disturbance as a result of cross-talk with the environment. A coordinated production of proteins can be envisioned to facilitate their interactions, the occurrence of which at the right place and right time could be essential for the information cascade. In contrast to proteins, genes including their copy numbers, positions on the linear DNA and 3-D chromatin are less variable and provide a more stable information storage. This study suggests that a coordinated and cellular-state dependent, highly regulated protein-protein interaction network can be achieved through usage of information stored in gene-gene interactions in 3-D chromatin structure. Such an information flow can lead to coordinated transcription (in time, and probably also in space) and eventually to functional protein-protein interactions. One can imagine that such protein-protein interactions involve not only pairs of proteins but also hetero-complexes formed by multiple proteins, the formation of which requires conceivably an even higher-level of coordination.

Hence, the stable information storage of genomic structure can also furnish intrinsic guidance on quantifications of tissue/cell-specific protein-protein interactions. In contrast to the fast accumulation of Hi-C data, high-throughput quantifications of tissue/cell-specific protein-protein interactions are still challenging. The genomic structure changes provide important knowledge and complementary information in predicting tissue-specific protein-protein interactions, which is expected to be of use in understanding the dynamic function of proteomics, as well as the resulted gene regulation network. In future studies, to understand the molecular mechanisms leading to the various molecular associations, we will thoroughly analyze the sequence and structure of proteins identified through the current chromatin structure analysis.

In summary, C_T_G is a consistent and robust framework for understanding the 3D genomic structure, by which we have detected a possible information flux of gene-gene interaction from genomic structure to transcriptional and translational levels. This study reveals important information of gene-gene physical interactions in 3D chromatin structure formation and their changes in cancer compared to normal tissues. We found that this physical contact information between genes at the DNA level is likely transferred to the protein level for at least a subset of genes. The underlying mechanisms and functions of the passage of genomic to transcription correlation and protein-protein interaction requires much more experimental and computational validations and tests. For a more decisive evaluation on the GGI and PPI relationship, concurrent measurement of them in the same cells at the single-cell level would be extremely valuable.

## Methods

### C_T_G algorithm

W denotes for the Hi-C contact map normalized by ICE. It is a positive symmetric matrix and is regarded as the adjacency matrix for a weighted connected network *G*(*V,E*), where the vertices *V=*{*v*_*1*_,*v*_*2*_,*…v*_*n*_} denote the non-overlapping genomic regions and the edges *E =* {*e*_*i,j*_} denote the contact strength between pairwise genomic regions. *D* is the diagonal degree matrix for the network, where the matrix element 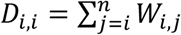. An 1-step transition probability matrix *P*^(*1*)^ can be derived by row-normalization of *W*:

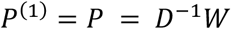

As *W* is diagonalizable, *P* is also diagonalizable:

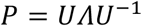

The eigenvectors U = {u_1_,u_2_,…,u_n_} reflect the characteristics of the reference matrix P. From the perspective of spectral analysis, the eigenvectors indicate the hierarchy of the network and the eigenvector corresponding to the largest eigenvalue indicates the most predominant hierarchy level of the network. Specific to genomic structures, the eigenvectors are respectively assigned to hierarchical structures, such as compartments and TAD structures. And the local systematic biases are more likely to be assigned to eigenvectors of small eigen values, as they are not global properties.

The k-step transition probability matrix *P*^(*k*)^ can be written as k-th power of *P*^(1)^:

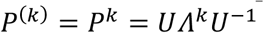

With step number *k* increasing, eigenvectors associated with genomic structure are preserved. Meanwhile, for a larger *k*, the contributing proportion of eigenvectors (corresponding eigenvalues) changes, where eigenvectors corresponding to larger eigenvalues of *Λ* gradually contribute more and *P*^(*k*)^ highlight predominant hierarchy level of the network. *P*^(*k*)^ converges to the invariant distribution quickly and the difference between *P*^(*k* − 1)^ and *P*^(*k*)^ decreases sharply, which means *P*^(*k*)^ provide less and less new information with k increasing. An exponential decay to was chosen to fit the convergence and a transition propensity matrix *S* within *k* steps is defined as:

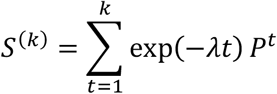

When *k* approaches infinity, *S*^*(k)*^ converge to M (Supplymentary note1):

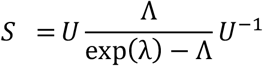

Therefore, the properties of *S* are independent from the value *k. S*_*i*_ denotes the *i*th row of *S* and represents the integrated diffusion propensity of the *i*th vertex. The L1 norm of *S*_*i*_ can be written as:

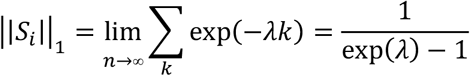

Considering the uniformity of the L1 norm of *S*_*i*_, we quantify the similarity between pairwise genomic regions *i* and *j* by calculating the L1 distance between *S*_*i*_ and *S*_*j*_. Hence, a C_T_G distance matrix is constructed from Hi-C contact matrix and the distance measures the similarity of pairwise genomic regions by their diffusion propensity in a genome-wide fashion.

### Hi-C experiment

#### Cell culture and fixation

HEK293 cells (American Type Culture Collection) were cultured at 37 °C under 5% CO_2_ in a humidified incubator. We cultured HEK293 cells in DMEM medium (Gibco, #11965092) with 10% fetal bovine serum and 1% penicillin–streptomycin. To gather the cells for Hi-C processing, the cells were washed twice using PBS, detached by adding 1 mL 0.25% trypsin-EDTA (Gibco, #25200056) to their culture dish, centrifuged at 500g for 5 min, and recovered in PBS buffer. The cells were counted by a cell counter to determine the concentration. For sample 0923-4, 1000 cells were extracted to a 0.5 mL Eppendorf Lobind Microcentrifuge tube (Eppendorf, #32119210) for each sample. For sample 1002-5 and 0923-2, 10,000 cells were extracted.

The cells were then fixed by adding formaldehyde (Sigma-Aldrich, #47608) to a final concentration of 2% at room temperature for 10 min then quenched by 0.2 M glycine (Sigma-Aldrich, #50046) for 10 min. The fixed cells were centrifuged at 2,500g for 5 min to discard the supernatant and washed by 0.5 mL PBS (Gibco, #20012027) once.

#### Hi-C processing

Hi-C experiments were performed followed methods described in Rao et al., 2014 with some modifications. Briefly, the fixed cell pallet was lysed in 100 µL Hi-C lysis The fixed cell pallet was lysed in 100 µL Hi-C lysis buffer (10 mM Tris-HCl pH=7.6 (Rockland, #MB-003), 10mM NaCl (Invitrogen AM9760G), 0.2% IGEPAL CA-720 (Sigma-Aldrich, #238589), 1x cOmplete protease inhibitor (Roche, #04693116001)) on ice for at least 30 min. The tubes were centrifuged to remove all the supernatant. 50 µL of 0.5% SDS (Invitrogen, #15553027) was added to each tube and incubate at 65°C for 20 min. To quench the reaction, 25 µL of 10% Triton X-100 (Sigma Aldrich, #T8787) was added and mix by pipetting up and down for several times. The tubes were then incubated at 37°C for 20 min. To perform chromatin digestion, 10 µL 10x NEBuffer2 (NEB, #B7002S), 10 µL 25U/µL MboI (NEB, #R0147L) and 5 µL water were added to each tube and incubate at 37°C with rotation for 24 h. MboI enzyme was inactivated at 62°C for 20 min. Fill-in mix which contains 14 µL 0.4 mM biotin-dATP (Invitrogen #19518018), 0.17 µL 10mM dTTP (NEB, #N0446S), 0.17 µL 10mM dGTP (NEB, #N0446S), 0.17 µL 10mM dCTP (NEB, #N0446S) and 3 µL 5U/µL DNA Polymerase I Large (Klenow) Fragment (NEB, #M0210V) was added and incubated at 37°C for 45min with rotation. Next, 12 µL 10% Triton X-100, 1.5 µL 100x BSA (NEB, #B9000S), 5 µL 10x T4 DNA ligase reaction buffer (NEB, #B0202S), 2 µL 400U/µL T4 DNA ligase (NEB, #M0202V), 10 µL 10mM ATP (NEB, #P0756S) and 2 µL water were added to each sample and the ligation reaction was carried out by incubating at room temperature with rotation for 24 h.

#### Library Construction

After ligation, DNA fragments were released by addition of 15 µL 10% SDS and 30 µL Proteinase K (Qiagen, #19133) to each tube followed by incubation at 50°C for 3h. The DNA fragments were purified by Ampure XP beads (volume ratio 1:1, Beckman Coulter, #A63881) and elute the DNA fragments in 27 µL water. Tagemtation was performed by adding 4 µL 8x TD buffer (80mM Tris-HCl pH=7.6, 40mM MgCl2 (Invitrogen, #AM9530G), 80% N,N Dimethylformamide (Sigma-Aldrich, #D4551)) and 1 µL TTE Mix V50 Tn5 enzyme (Vazyme, #TD501) to the 27 µL DNA template. The tubes were incubated at 55°C for 1h. To stop the reation, 8 µL 5x TS (Vazyme, #TD503) was added to each tube and incubate at room temperature for 5 min. To prepare Dynabeads M-280 streptavidin (Invitrogen, #11206D) for the capture of ligation junctions, 25 µL streptavidin beads was washed by 1x BW buffer (5mM Tris-HCl pH=7.6, 0.5mM EDTA (Invitrogen #AM9260G), 1M NaCl) and resuspended in 13 µL 4x BW buffer (20mM Tris-HCl pH=7.6, 2mM EDTA, 4M NaCl) for each sample. The beads were then mixed with 40 µL tagmentation mix and incubate at room temperature overnight with rotation. The streptavidin beads were washed twice with 1x BW buffer, twice with 10mM Tris-HCl pH=7.6 and resuspended in 20 µL 10mM Tris-HCl pH=7.6. PCR amplification was carried out by addition of 5 µL 10 µM Nextera index mix(Vazyme, #TD203) and 25 µL Q5 High-Fidelity 2X master mix (NEB, #M0492S) to the 20 µL sample. PCR program was set as follows:

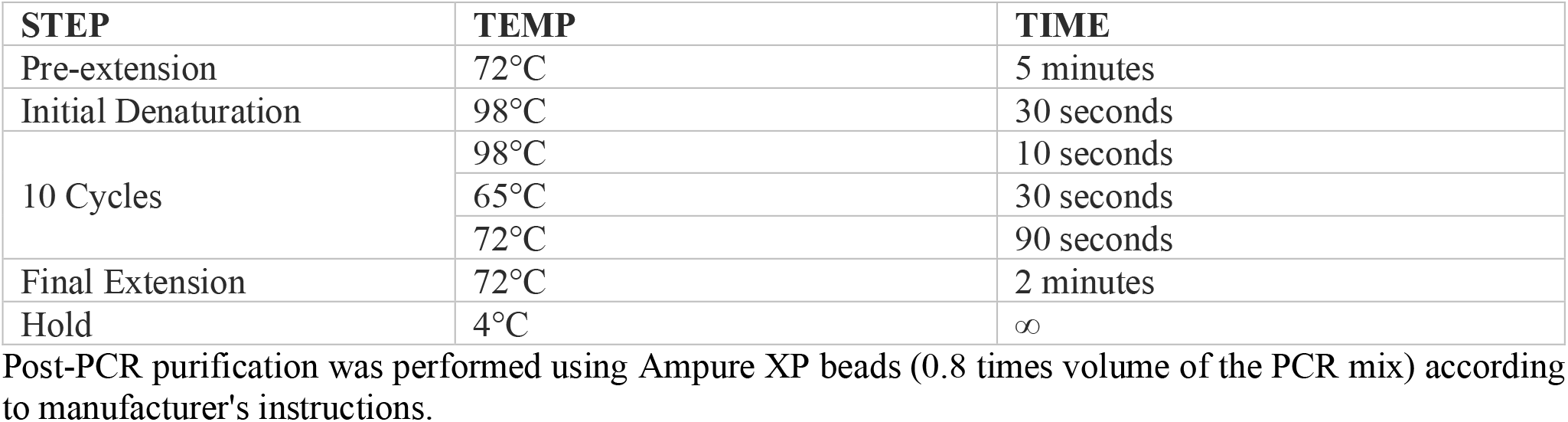

| STEP | TEMP | TIME |
| --- | --- | --- |
| Pre-extension | 72°C | 5 minutes |
| Initial Denaturation | 98°C | 30 seconds |
| 10 Cycles | 98°C | 10 seconds |
|  | 65°C | 30 seconds |
|  | 72°C | 90 seconds |
| Final Extension | 72°C | 2 minutes |
| Hold | 4°C | ∞ |

Post-PCR purification was performed using Ampure XP beads (0.8 times volume of the PCR mix) according to manufacturer's instructions.

#### Library QC and sequencing

The libraries were quantified using Qubit 1x dsDNA HS Assay kits (Invitrogen, #Q33230) and the size distribution was assessed using 5200 Fragment Analyzer System (Agilent, #M5310AA). The qualified libraries were then quantified by qPCR and sequenced by 2x 150 bp paired-end run on a Novaseq 6000 System (Illumina).

#### Sequence Processing

Paired-end reads were first under adaptor trimming using Cutadapt (Martin, 2011, version 2.10) with default arguments. Reads shorter than 20 bp were filtered out after adapter trimming. Trimmed reads were mapped to Genome Reference Consortium Human Build 37 (hg19, downloaded from UCSC, https://hgdownload.soe.ucsc.edu/goldenPath/hg19/bigZips) and processed by HiC-Pro (Servant et al., 2015, version 2.11.4) using default settings. The contact matrix extracted by HiC-Pro were then used in downstream analysis.

### Hi-C data analysis

For normal and tumor colon samples, we used Hi-C data from GSE133928, the normal samples are BRD3162N-sb, BRD3179N, BRD3187N, BRD3462N, the tumor samples are BRD3162, BRD3179, BRD3187, BRD3146. For HCT116 cell line, we used Hi-C data from GSE133928. For MCF-7 and MCF-10A cell line, we used the samples GSE165570. Hi-C matrices were normalized using the ICE algorithm (Imakaev, Maxim; Fundenberg, Geoffrey; Patton McCord, Rachel; Naumova, Natalia; Goloborodko, Anton; Lajoie, Bryan R.; Dekker, Job; Mirny, 2012).

### Contrast ratio

Sobel operator is a discrete derivative operator for edge detection which is defined as:

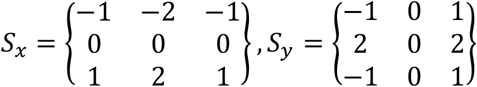

Convolution was performed on a distance map I with the Sobel operator as the kernel:

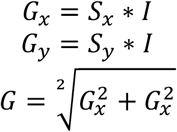

Distinct edges will be emphasized by *G* for a distance map with ‘chess-like squares’. Therefore, G reflects the contrast ratio of the genomic “squares” with distinct edges over their proximal neighbors and the mean of G is defined as the overall contrast ratio of the distance map.

### Laplacian Eigenmaps

Given a C_T_G distance map *I*, it’s transformed into weight matrix *W* by a exponential kernal :

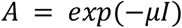

*µ* reflects the scale of genomic structure we focused on. A large *µ* amplifies weights of short-distance and a small *µ* amplifies weights of long-distance. To avoid the impact uneven degree distribution, the normalized Laplacian *L* is constructed:

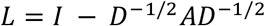

Where *D* is the degree matrix for *W*.

*L* is diagonalizable and the bottom 3 eigenvectors E0, E1,E2 is computed. E0 is excluded as it is not informative:

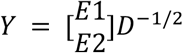

The coordinates for n genomic regions {y_1_,|y_2_| … |y_n_} ∈ ℝ^2^ is acquired by converting the columns of *Y* into 2-dimensional vectors:

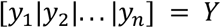

### Gene expression data

We downloaded all available tumor-normal pairwise somatic expression data for patients from TCGA GDC Data Portal (https://portal.gdc.cancer.gov) and selected expression data with more than 10 patients for 17 cancer types/subtypes. All expression data were converted to TPM (transcripts per million) format.

### Protein-Protein interaction data

To build a comprehensive protein-protein interactome, we assembled protein-protein interactions from 3 sources: (1) PPIs from STRING project (https://www.string.com), (2) HCT116-related PPIs from BioPlex project, (3) MCF-10A-related and MCF-7-related PPIs from Kim, M. *et al*. The cell-specific PPIs were determined from (2)(3) with PPI score ≥ 0.65 and ≥8-fold change.

### Genomic neighborhood and csGGIs

The neighborhood for a given genomic region is defined by its radial distribution function(RDF), taken a small proportion of genomic regions within the neighborhood. The diameter of the neighborhood is determined by boundary of the highest characteristic peak, where the slope of tangent line of the cumulative RDF is calculated, and tangent line with lagerst slope is chosen as a guideline. The diameter is quantified by the point that the guidline intersects with the x axis. The neighborhood for a given genomic region is then settled. Pairwise genomic regions within each other’s neighborhood is defined to have gene-gene interactions (GGIs).

The fold-change of C_T_G distance between tumor samples and normal samples is calculated, where *m* denotes for the mean of fold-change and *σ* denotes for the standard deviation. The cancer specific GGIs (csGGIs) are GGIs from tumor samples with extreme change in C_T_G distance (fold-change < *m* − 3*σ*, according to 3*σ* rule).

### Gene function analysis

GO enrichment analysis of all the given gene clusters in this work was conducted using DAVID (https://david.ncifcrf.gov). Individual gene functions were obtained from GeneCards (https://www.genecards.org). Cancer genes were obtained from COSMIC (https://cancer.sanger.ac.uk/cosmic).

### Visualization

The PyMOL program (Schrodinger LLC, 2015) was employed to visualize the genomic structure (Xie et al., 2017) in Figure 1. The VMD program (Humphrey et al., 1996) was employed to render the protein structure.

## Supplementary Materials

Figure S1: 2D Laplacian eigenmap of colon cancer, Figure S2: Correspondence of gene-gene proximity and RNA co-regulation in leukemia, Figure S3: Subgraph of csGGI network with ERBB3, Figure S4: Subgraph of csGGI network with HRAS, Figure S5: Subgraph of csGGI network with PTK6., Figure S6: Correspondance of gene-gene proximity and STRING PPIs in normal colon sample, Figure S7: The distribution of RNA correlation with STRING PPI for 17 cancer types, Figure S8: The distribution of RNA correlation for colon cancer, Table S1: Function enrichment analysis, Table S1: Functional annotation clustering of colon csGGIs, Table S2: Functional annotation clustering of HCT116 structural-related intra-chromosomal PPIs; Table S3: Functional annotation clustering of HCT116 structural-related inter-chromosomal PPIs.

## Author Contributions

Conceptualization, YQ.G.; Data curation, YY.He, Y.X.; Formal analysis, YY.He, Y.X., YP.H.; Experiment, L.L.; Funding acquisition, YQ.G.; Investigation, YY.He, Y.X., YP.H., JY.W. and YQ.G.; Supervision, YQ.G.; Visualization, YY.He and YP.H.; Writing – original draft, YY.He; Writing – review & editing, YY.Huang and YQ.G.

## Data Availability Statement

All data analyzed during this study are publicly available. The detailed data accession can be found in Methods section. <u>Liulu</u>

## Acknowledgments

The results shown here are part based upon data generated by the TCGA Research Network (http://cancergenome.nih.gov).

## Supplementary note

Proof 1: eigenvalues Λ of the P are within range of [-1,1].

For any eigenvector X of P:

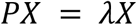

The maximum element of X is denoted as *x*_*max*_, and the minimum element of X is denoted as *x*_*min*_, As the row summation of P is normalized to 1, and P is positive,

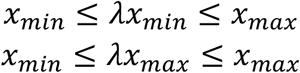

Therefore,

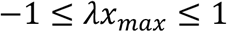

Proof 2: When n approaches infinity, the transition propensity matrix M^(n)^ is convergent.

P is diagonalizable:

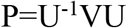

P^(k)^ can be written as:

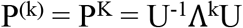

S^(n)^ is the weighted summation of P^(k)^:

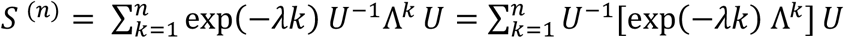

According to associative law of multiplication :

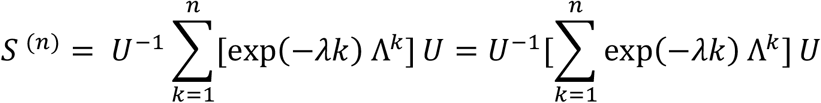

When *n* approaches infinity, we have

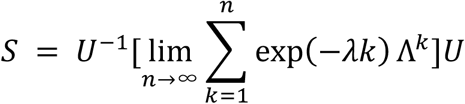

In the above equation, exp(−*λk*) Λ^*k*^ is a geometric progression, and

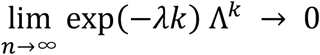

Therefore, the summation over exp(−*λk*) Λ^*k*^ is convergent, and

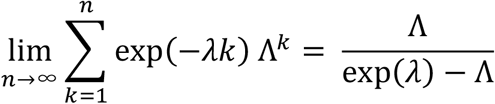

S is then also convergent and

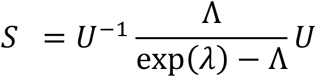

**Figure S1.**
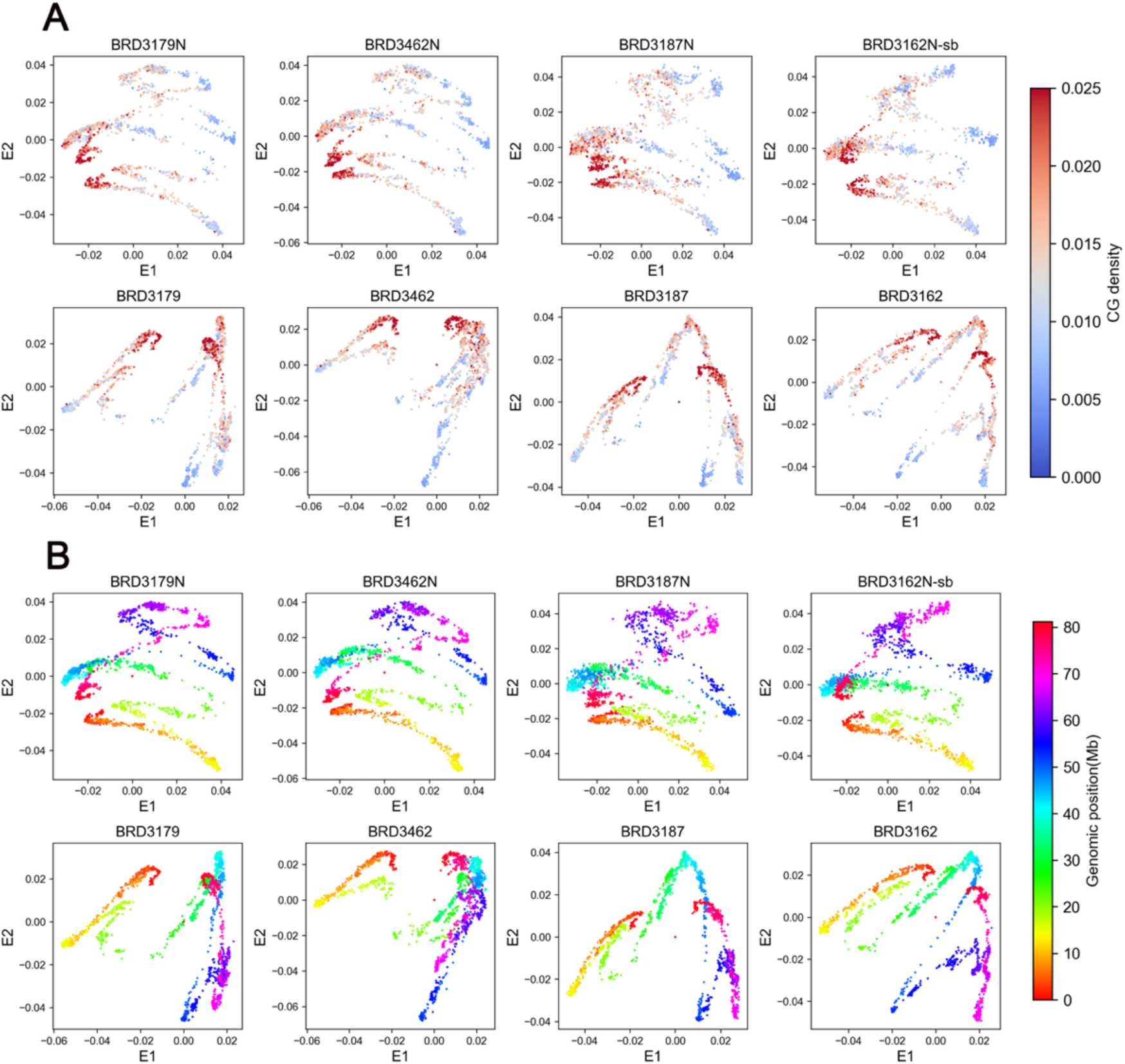
2D Laplacian eigenmap of colon cancer. **(A)** The 2D Laplacian Eigenmaps of C_T_G distance matrices for pairwise normal (upper pannel) and tumor (lower pannel) colon samples. Each point represents a 40kb genomic region. The color is used to represent the CpG density of the corresponding genomic region. **(B)** The color is used to represent the genomic position of the corresponding genomic region.

**Figure S2.**
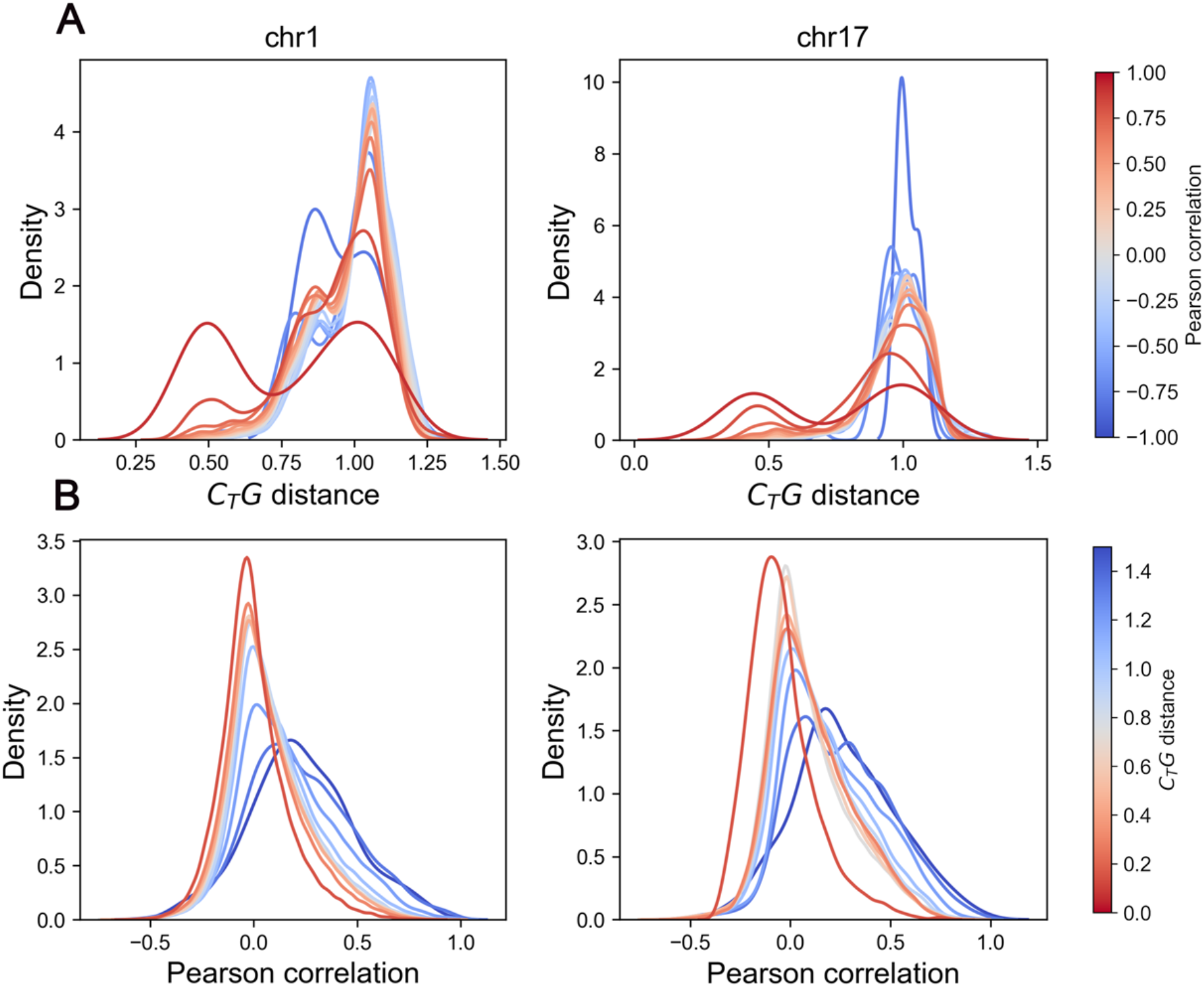
Correspondence of gene-gene proximity and RNA co-regulation in leukemia. (**A**) The distribution of transcriptional Pearson correlation under different C_T_G distance of chromosome 1(left) and chromosome 17(right), the color of each line indictes corresponding C_T_G distance. (**B**) The distribution of C_T_G distance under different Pearson correlation of chromosome 1(left) and chromosome 17(right), the color of each line indictes corresponding Pearson correlation coefficient.

**Figure S3.**
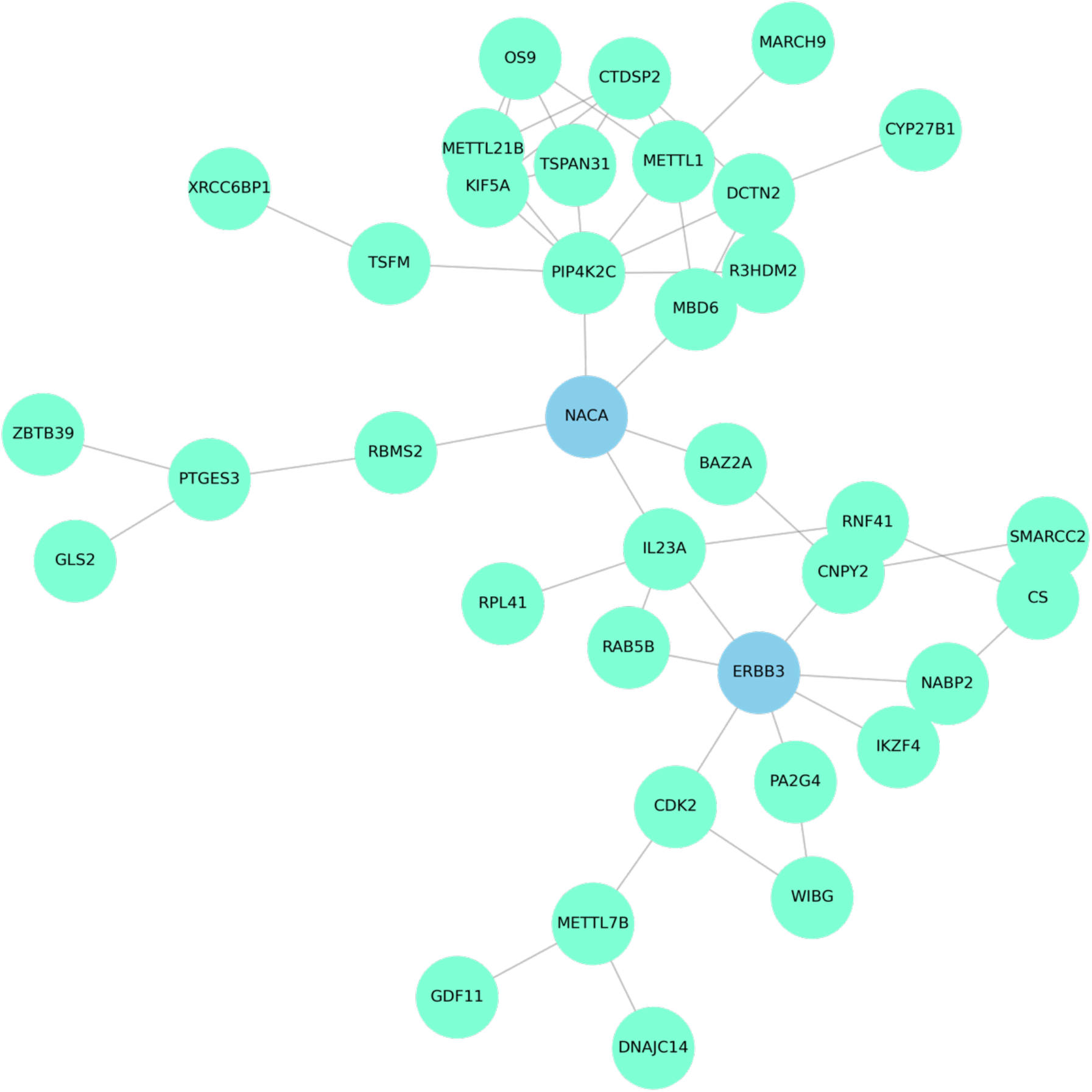
Subgraph of csGGI network with ERBB3.

**Figure S4.**
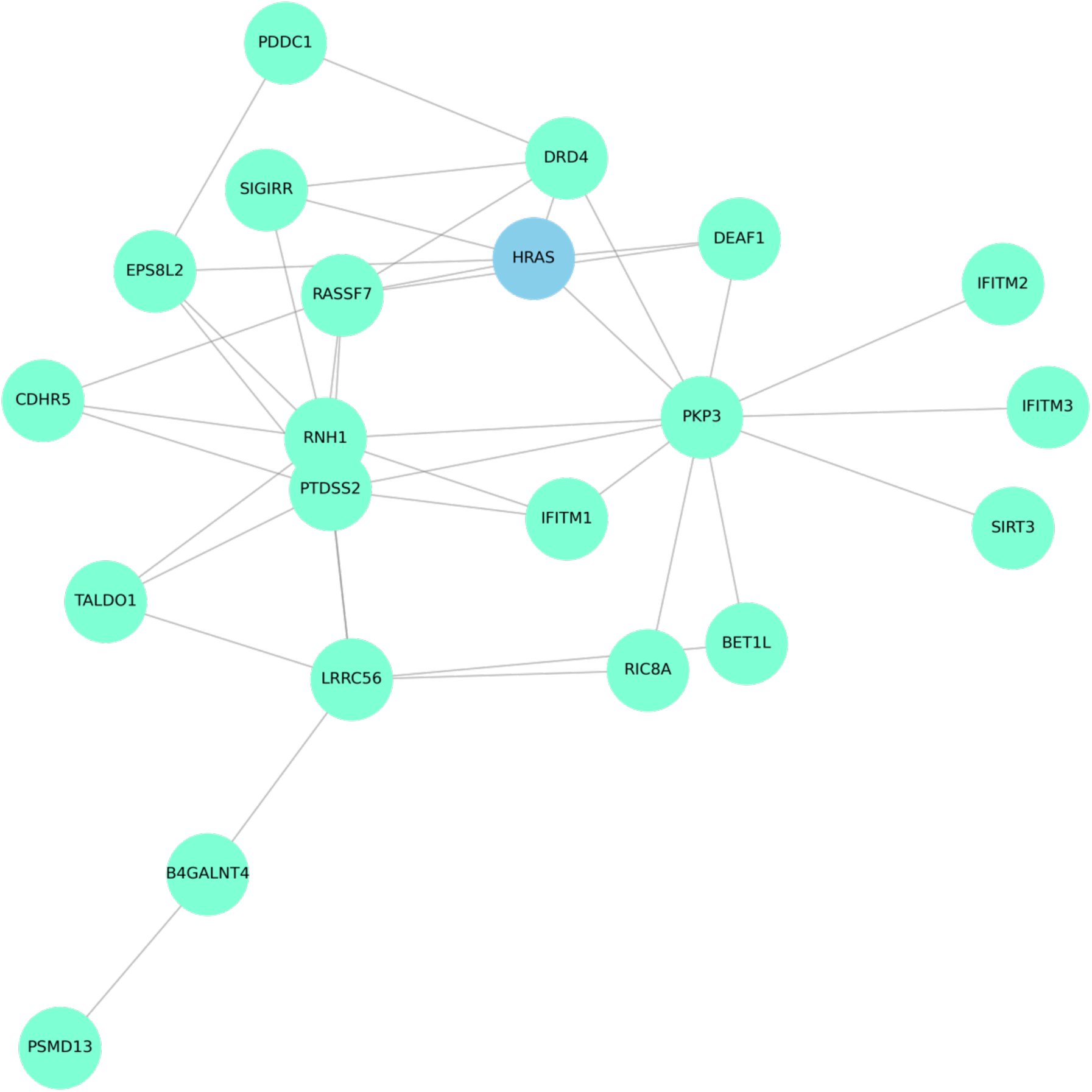
Subgraph of csGGI network with HRAS.

**Figure S5.**
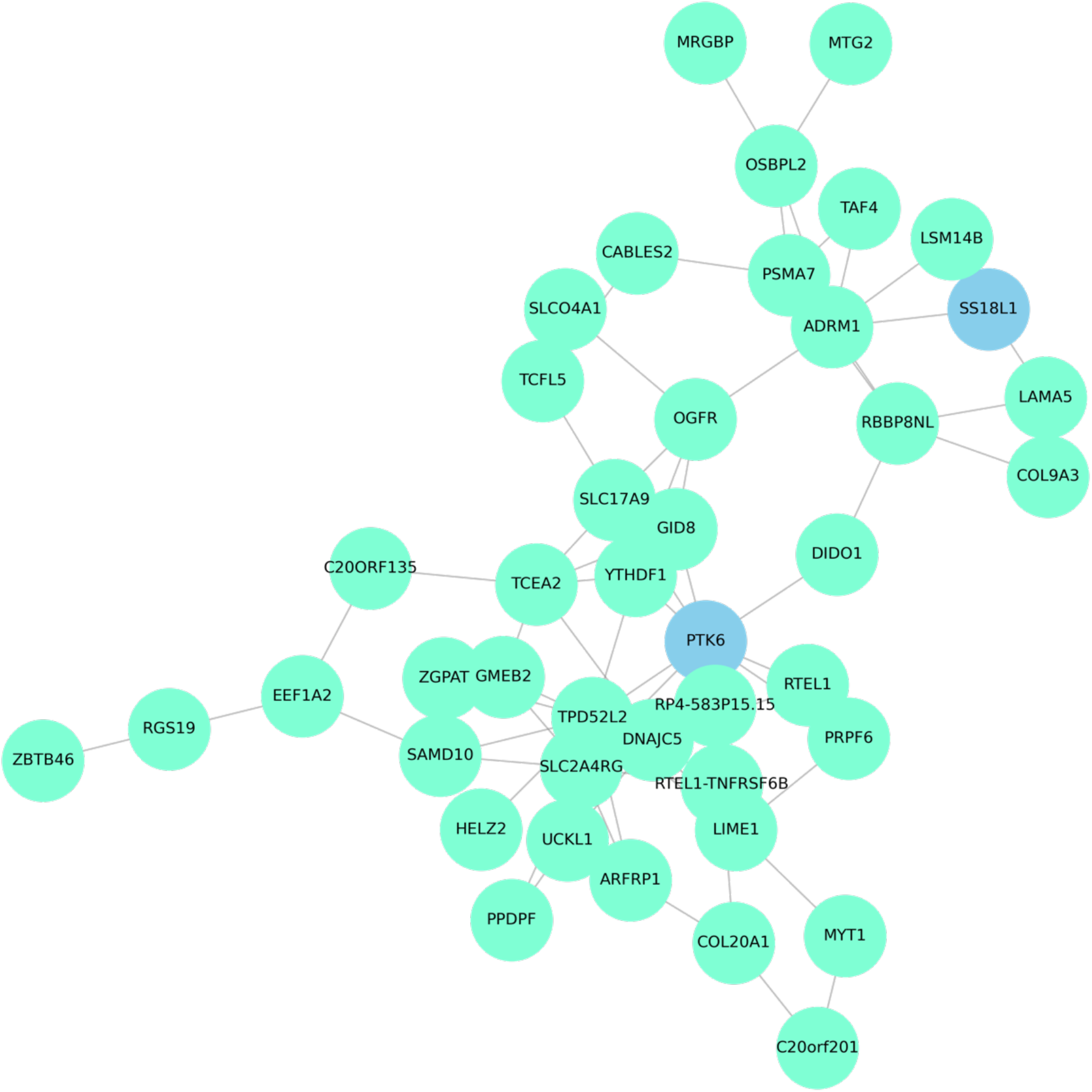
Subgraph of csGGI network with PTK6.

**Figure S6.**
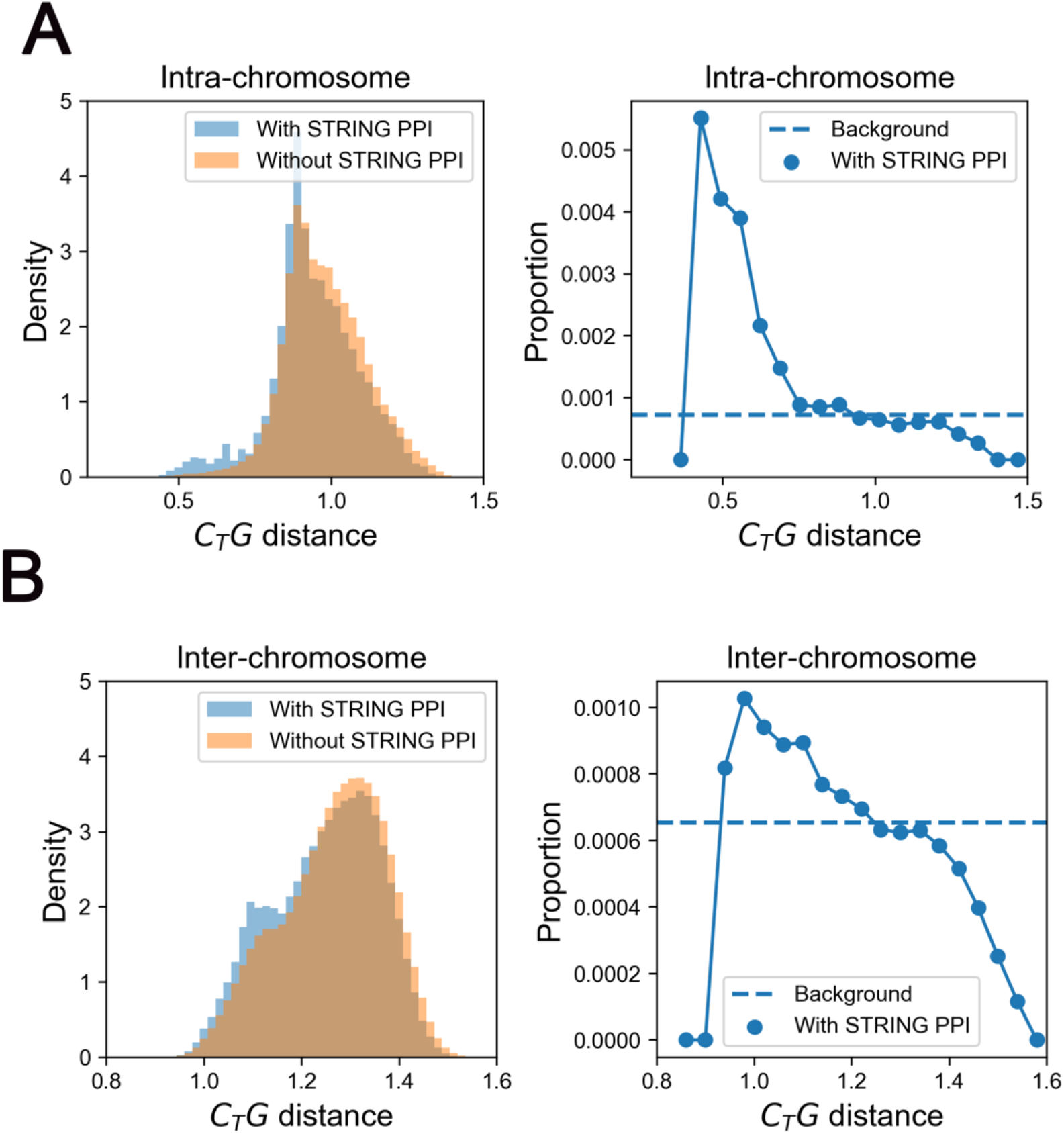
Correspondance of gene-gene proximity and STRING PPIs in normal colon sample. (**A**) The distribution of C_T_G distance between intra-chromosomal gene pairs with and without STRING PPIs for whole chromosome in tumor sample (left pannel); The proportion of intra-chromosomal gene-pairs with STRING PPI at different C_T_G distances in normal sample (right pannel). (**B**) The distribution of C_T_G distance between inter-chromosomal gene pairs with and without STRING PPIs for whole chromosome in normal sample (left pannel); The proportion of inter-chromosomal gene-pairs with STRING PPI at different C_T_G distances in normal sample (right pannel).

**Figure S7.**
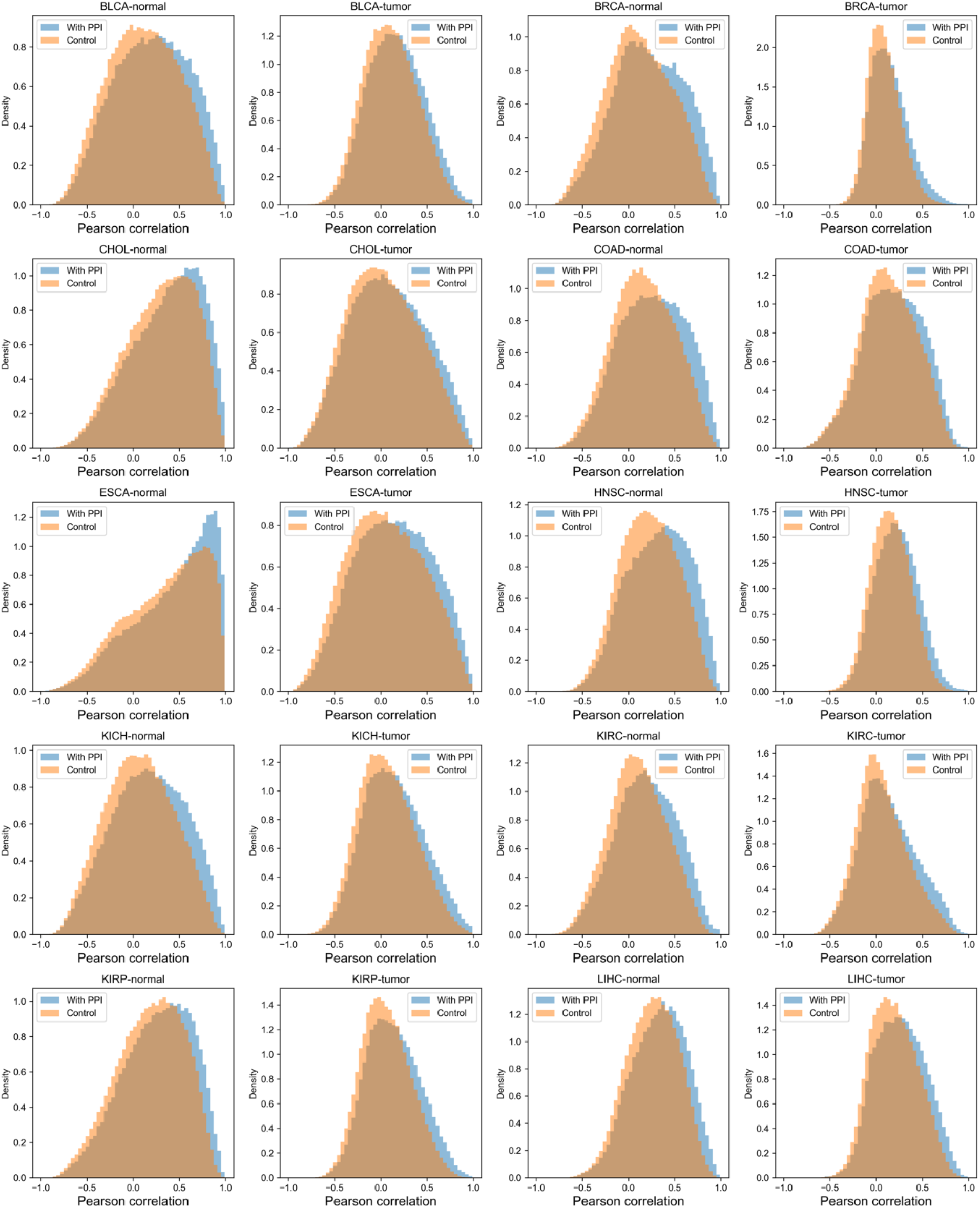
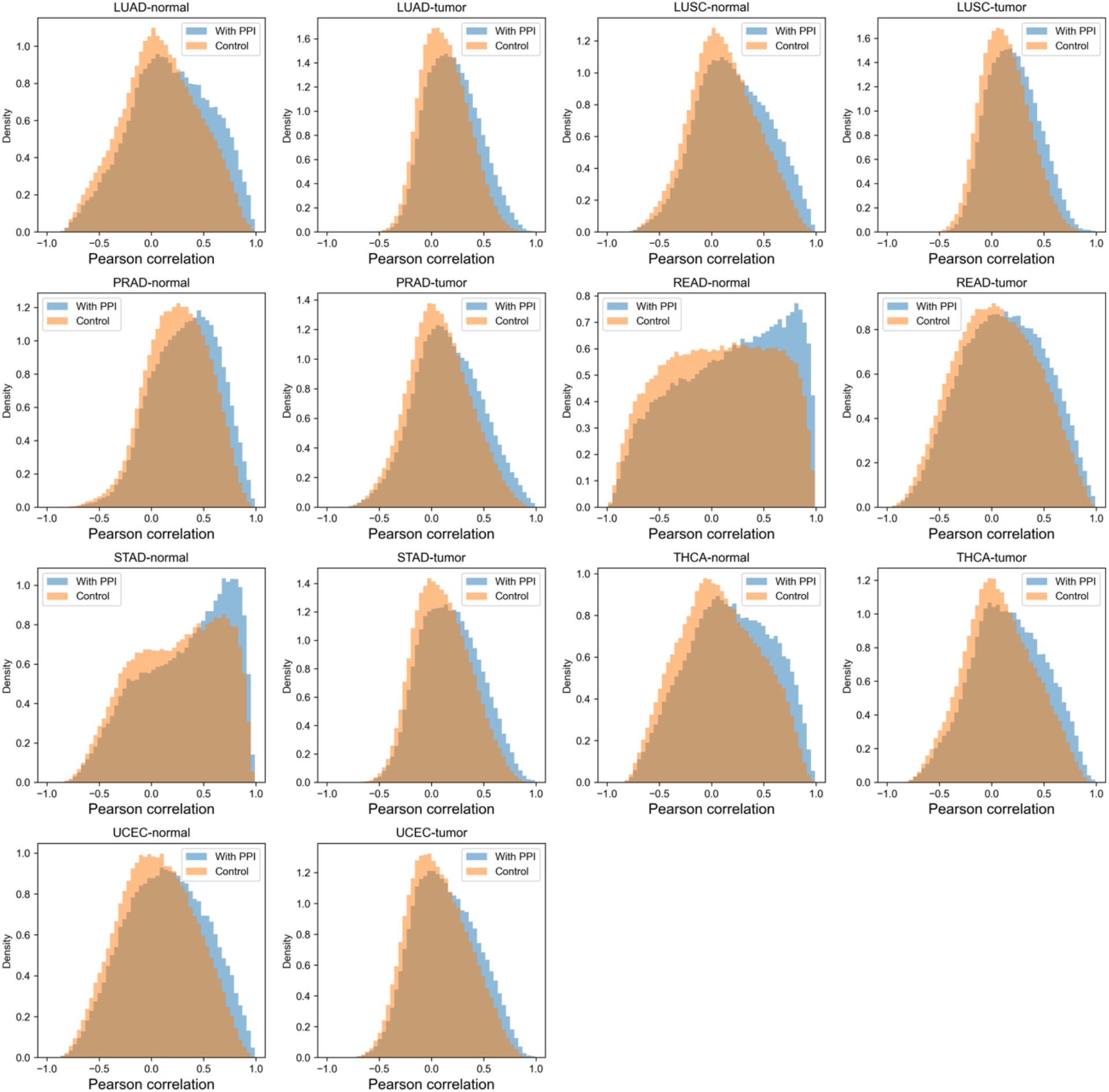
The distribution of RNA correlation with STRING PPI for 17 cancer types. The distributions of gene-pairs with STRING PPI are colored blue and the control groups are colored orange. All samples show similar patterns that gene-pairs with STRING PPI are more correlated in transcriptional level.

**Figure S8.**
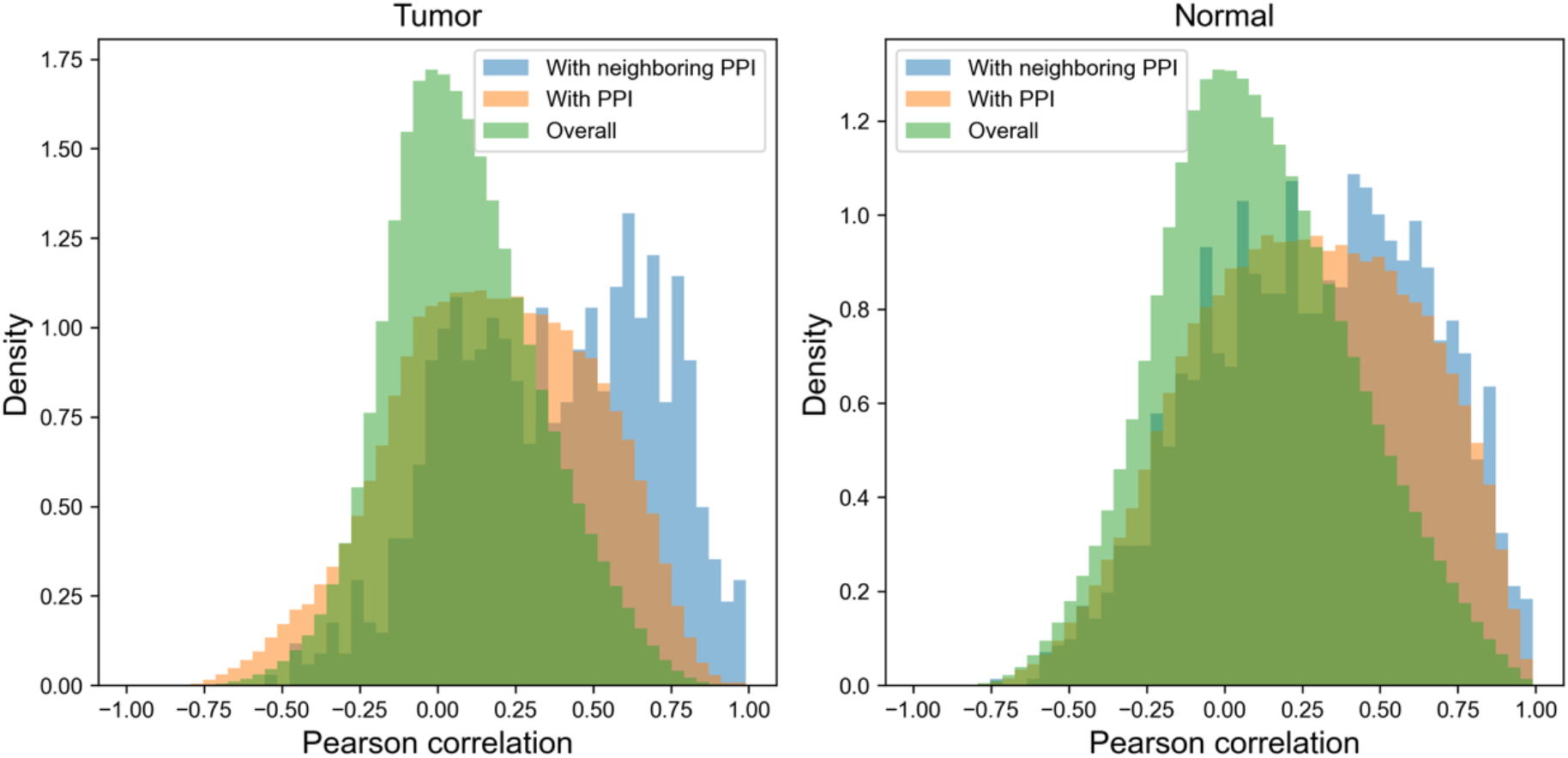
The distribution of RNA correlation for colon cancer. Gene-pairs with both STRING PPI and GGIs are colored blue and gene-pairs with only STRING PPI are colored orange and the control group is colored green. Gene-pairs with both STRING PPI and GGIs are more correlated in transcriptional level for both tumor and normal colon sample.

**Table S1.** Functional annotation clustering of colon csGGIs.

| Annotation Cluster 1: Enrichment Score: 1.5327833580397092 |  |  |  |
| --- | --- | --- | --- |
| Term | Count | PValue | Benjamini |
| hsa01521: EGFR tyrosine kinase inhibitor resistance | 7 | 4.63E-05 | 0.00920864 |
| h_pyk2Pathway: Links between Pyk2 and Map Kinases | 5 | 8.51E-05 | 0.00802485 |
| h_at1rPathway: Angiotensin II mediated activation of JNK Pathway via Pyk2 dependent signaling | 5 | 1.65E-04 | 0.00802485 |
| hsa05219: Bladder cancer | 5 | 3.55E-04 | 0.03383294 |
| h_malPathway: Role of MAL in Rho-Mediated Activation of SRF | 4 | 5.67E-04 | 0.01608635 |
| hsa04662: B cell receptor signaling pathway | 6 | 5.77E-04 | 0.03383294 |
| hsa04012: ErbB signaling pathway | 6 | 6.80E-04 | 0.03383294 |
| h_rasPathway: Ras Signaling Pathway | 4 | 0.00101341 | 0.0196602 |
| hsa04140: Autophagy - animal | 7 | 0.00108928 | 0.03764245 |
| hsa05231: Choline metabolism in cancer | 6 | 0.00129541 | 0.03764245 |
| hsa04370: VEGF signaling pathway | 5 | 0.00142489 | 0.03764245 |
| hsa05205: Proteoglycans in cancer | 8 | 0.00151326 | 0.03764245 |
| h_sam68Pathway: Regulation of Splicing through Sam68 | 3 | 0.00163092 | 0.02636658 |
| GO: 2000641~regulation of early endosome to late endosome transport | 3 | 0.0017761 | 0.66330982 |
| hsa04810: Regulation of actin cytoskeleton | 8 | 0.00215053 | 0.04444882 |
| h_erkPathway: Erk1/Erk2 Mapk Signaling pathway | 4 | 0.00223416 | 0.03095906 |
| hsa04664: Fc epsilon RI signaling pathway | 5 | 0.00241051 | 0.04444882 |
| hsa05211: Renal cell carcinoma | 5 | 0.00254296 | 0.04444882 |
| hsa04917: Prolactin signaling pathway | 5 | 0.00268033 | 0.04444882 |
| hsa05223: Non-small cell lung cancer | 5 | 0.00297021 | 0.04546711 |
| hsa05220: Chronic myeloid leukemia | 5 | 0.00361302 | 0.05135645 |
| h_metPathway: Signaling of Hepatocyte Growth Factor Receptor | 4 | 0.00379947 | 0.03988317 |
| hsa04650: Natural killer cell mediated cytotoxicity | 6 | 0.00389495 | 0.05167298 |
| h_fmlpPathway: fMLP induced chemokine gene expression in HMC-1 cells | 4 | 0.00411167 | 0.03988317 |
| h_integrinPathway: Integrin Signaling Pathway | 4 | 0.00411167 | 0.03988317 |
| hsa05210: Colorectal cancer | 5 | 0.00561992 | 0.06989776 |
| h_cdk5Pathway: Phosphorylation of MEK1 by cdk5/p35 down regulates the MAP kinase pathway | 3 | 0.00586627 | 0.05172985 |
| hsa04540: Gap junction | 5 | 0.00609515 | 0.07134909 |
| Annotation Cluster 2: Enrichment Score: 1.5241353790689698 |  |  |  |
| Term | Count | PValue | Benjamini |
| GO: 0039702~viral budding via host ESCRT complex | 3 | 0.01072312 | 1 |
| GO: 0036258~multivesicular body assembly | 3 | 0.02069957 | 1 |
| GO: 0043162~ubiquitin-dependent protein catabolic process via the multivesicular body sorting pathway | 3 | 0.02069957 | 1 |
| GO: 0090148~membrane fission | 3 | 0.03330408 | 1 |
| Annotation Cluster 3: Enrichment Score: 1.453480544316328 |  |  |  |
| Term | Count | PValue | Benjamini |
| KW-0653~Protein transport | 12 | 0.01295629 | 0.24467061 |
| GO: 0015031~protein transport | 8 | 0.0389436 | 1 |
| KW-0967~Endosome | 9 | 0.08642541 | 0.79943504 |
| Annotation Cluster 4: Enrichment Score: 1.444194122407839 |  |  |  |
| Term | Count | PValue | Benjamini |
| KW-0648~Protein biosynthesis | 6 | 0.00775783 | 0.24467061 |
| KW-0396~Initiation factor | 3 | 0.07027195 | 0.71901399 |
| GO: 0003743~translation initiation factor activity | 3 | 0.08528948 | 1 |
| Annotation Cluster 5: Enrichment Score: 1.3130484622630447 |  |  |  |
| Term | Count | PValue | Benjamini |
| GO: 0051402~neuron apoptotic process | 5 | 0.001971 | 0.66330982 |
| GO: 0043524~negative regulation of neuron apoptotic process | 4 | 0.10479492 | 1 |
| Annotation Cluster 6: Enrichment Score: 1.2661570708970376 |  |  |  |

| Term | Count | PValue | Benjamini |
| --- | --- | --- | --- |
| GO: 0097542~ciliary tip | 4 | 0.00441052 | 0.19343661 |
| GO: 0005813~centrosome | 11 | 0.00465311 | 0.19343661 |
| GO: 0042073~intraciliary transport | 3 | 0.02328684 | 1 |
| GO: 0005929~cilium | 6 | 0.02622916 | 0.44898151 |
| Annotation Cluster 7: Enrichment Score: 1.2170182899365116 |  |  |  |
| Term | Count | PValue | Benjamini |
| GO: 0019216~regulation of lipid metabolic process | 4 | 0.00293104 | 0.66330982 |
| GO: 0042752~regulation of circadian rhythm | 3 | 0.08129563 | 1 |

**Table S2.** Functional annotation clustering of HCT116 structural-related intra-chromosomal PPIs.

| Annotation Cluster 1: Enrichment Score: 18.886710115925517 |  |  |
| --- | --- | --- |
| Term | PValue | Benjamini |
| GO: 0007156~homophilic cell adhesion via plasma membrane adhesion molecules | 7.08E-39 | 1.38E-35 |
| GO: 0007399~nervous system development | 5.72E-17 | 5.59E-14 |
| GO: 0007155~cell adhesion | 1.66E-15 | 1.08E-12 |
| GO: 0005509~calcium ion binding | 1.07E-14 | 6.82E-12 |
| GO: 0005887~integral component of plasma membrane | 5.09E-12 | 2.37E-09 |
| Annotation Cluster 2: Enrichment Score: 5.486720966179876 |  |  |
| Term | PValue | Benjamini |
| hsa05320: Autoimmune thyroid disease | 8.92E-19 | 2.45E-16 |
| GO: 0002323~natural killer cell activation involved in immune response | 1.09E-10 | 5.32E-08 |
| hsa05169: Epstein-Barr virus infection | 4.37E-10 | 6.01E-08 |
| GO: 0005132~type I interferon receptor binding | 2.03E-09 | 6.44E-07 |
| hsa05152: Tuberculosis | 8.01E-09 | 3.96E-07 |
| GO: 0033141~positive regulation of peptidyl-serine phosphorylation of STAT protein | 1.02E-08 | 3.98E-06 |
| hsa05164: Influenza A | 1.78E-08 | 5.91E-07 |
| hsa05163: Human cytomegalovirus infection | 1.93E-08 | 5.91E-07 |
| hsa05168: Herpes simplex virus 1 infection | 5.62E-08 | 1.54E-06 |
| GO: 0002286~T cell activation involved in immune response | 7.08E-08 | 2.18E-05 |
| GO: 0006959~humoral immune response | 1.07E-07 | 2.61E-05 |
| GO: 0019221~cytokine-mediated signaling pathway | 5.31E-07 | 1.01E-04 |
| hsa04623: Cytosolic DNA-sensing pathway | 5.84E-07 | 1.46E-05 |
| GO: 0042100~B cell proliferation | 6.20E-07 | 1.01E-04 |
| GO: 0060337~type I interferon signaling pathway | 6.20E-07 | 1.01E-04 |

| hsa05170: Human immunodeficiency virus 1 infection | 6.38E-07 | 1.46E-05 |
| --- | --- | --- |
| hsa05167: Kaposi sarcoma-associated herpesvirus infection | 6.93E-07 | 1.47E-05 |
| GO: 0043330~response to exogenous dsRNA | 7.02E-07 | 1.05E-04 |
| hsa05165: Human papillomavirus infection | 1.66E-06 | 3.26E-05 |
| hsa04650: Natural killer cell mediated cytotoxicity | 4.93E-06 | 8.47E-05 |
| hsa05162: Measles | 1.55E-05 | 2.37E-04 |
| GO: 0051607~defense response to virus | 2.19E-05 | 0.00237571 |
| GO: 0030183~B cell differentiation | 2.44E-05 | 0.0025043 |
| hsa04622: RIG-I-like receptor signaling pathway | 8.08E-05 | 0.00105761 |
| hsa05161: Hepatitis B | 8.57E-05 | 0.00107156 |
| GO: 0002250~adaptive immune response | 1.05E-04 | 0.00972708 |
| hsa05200: Pathways in cancer | 1.24E-04 | 0.00148135 |
| GO: 0098586~cellular response to virus | 1.42E-04 | 0.01258975 |
| hsa04217: Necroptosis | 2.67E-04 | 0.00271968 |
| hsa04620: Toll-like receptor signaling pathway | 3.80E-04 | 0.00373633 |
| hsa05417: Lipid and atherosclerosis | 5.11E-04 | 0.00484296 |
| hsa05160: Hepatitis C | 8.40E-04 | 0.00745575 |
| hsa04936: Alcoholic liver disease | 0.00123367 | 0.01028057 |
| GO: 0005125~cytokine activity | 0.00232746 | 0.21113399 |
| hsa04630: JAK-STAT signaling pathway | 0.00350412 | 0.02604415 |
| hsa04060: Cytokine-cytokine receptor interaction | 0.00466886 | 0.03292144 |
| hsa04151: PI3K-Akt signaling pathway | 0.00537873 | 0.0369788 |
| hsa05171: Coronavirus disease - COVID-19 | 0.00802705 | 0.05133577 |
| hsa04621: NOD-like receptor signaling pathway | 0.00900205 | 0.05626278 |
| GO: 0005126~cytokine receptor binding | 0.01312996 | 0.75795707 |
| Annotation Cluster 3: Enrichment Score: 4.299091183648573 |  |  |
| Term | PValue | Benjamini |
| hsa05320: Autoimmune thyroid disease | 8.92E-19 | 2.45E-16 |
| GO: 0071556~integral component of lumenal side of endoplasmic reticulum membrane | 3.08E-10 | 7.16E-08 |
| hsa05330: Allograft rejection | 2.03E-09 | 1.86E-07 |
| hsa05332: Graft-versus-host disease | 6.58E-09 | 3.96E-07 |
| hsa04940: Type I diabetes mellitus | 8.64E-09 | 3.96E-07 |
| GO: 0042605~peptide antigen binding | 1.09E-08 | 2.30E-06 |
| hsa04612: Antigen processing and presentation | 1.11E-08 | 4.37E-07 |
| GO: 0042613~MHC class II protein complex | 5.37E-08 | 8.32E-06 |
| GO: 0019882~antigen processing and presentation | 7.82E-08 | 2.18E-05 |
| GO: 0002504~antigen processing and presentation of peptide or polysaccharide antigen via MHC class II | 1.75E-07 | 3.79E-05 |
| GO: 0032395~MHC class II receptor activity | 2.70E-07 | 4.28E-05 |
| GO: 0002503~peptide antigen assembly with MHC class II protein complex | 8.60E-07 | 1.20E-04 |
| GO: 0002381~immunoglobulin production involved in immunoglobulin mediated immune response | 1.30E-06 | 1.70E-04 |
| hsa05416: Viral myocarditis | 3.00E-06 | 5.50E-05 |
| GO: 0012507~ER to Golgi transport vesicle membrane | 3.18E-06 | 2.95E-04 |
| GO: 0019886~antigen processing and presentation of exogenous peptide antigen via MHC class II | 3.77E-06 | 4.60E-04 |
| hsa04145: Phagosome | 9.83E-06 | 1.59E-04 |
| hsa05166: Human T-cell leukemia virus 1 infection | 1.99E-05 | 2.87E-04 |
| GO: 0023026~MHC class II protein complex binding | 3.50E-05 | 0.00444635 |
| hsa05150: Staphylococcus aureus infection | 3.91E-05 | 5.38E-04 |
| GO: 0050870~positive regulation of T cell activation | 8.58E-05 | 0.00838156 |
| hsa05310: Asthma | 1.29E-04 | 0.00148193 |
| GO: 0030666~endocytic vesicle membrane | 1.55E-04 | 0.01200662 |
| hsa05145: Toxoplasmosis | 1.61E-04 | 0.00177432 |
| hsa05140: Leishmaniasis | 1.71E-04 | 0.00180795 |
| GO: 0005765~lysosomal membrane | 2.22E-04 | 0.0147493 |
| GO: 0006955~immune response | 3.83E-04 | 0.03118785 |
| GO: 0002486~antigen processing and presentation of endogenous peptide antigen via MHC class I via ER pathway, TAP-independent | 5.93E-04 | 0.046337 |
| GO: 0042612~MHC class I protein complex | 7.87E-04 | 0.04053694 |
| hsa04514: Cell adhesion molecules | 8.40E-04 | 0.00745575 |
| GO: 0000139~Golgi membrane | 0.00109335 | 0.04621875 |
| hsa04640: Hematopoietic cell lineage | 0.00111535 | 0.00958507 |
| hsa05321: Inflammatory bowel disease | 0.00148522 | 0.01201284 |
| hsa04672: Intestinal immune network for IgA production | 0.00165846 | 0.01303076 |
| GO: 0030658~transport vesicle membrane | 0.00190334 | 0.06808101 |
| hsa05322: Systemic lupus erythematosus | 0.00299623 | 0.02288785 |
| GO: 0001916~positive regulation of T cell mediated cytotoxicity | 0.00301056 | 0.22613968 |
| GO: 0030670~phagocytic vesicle membrane | 0.00555792 | 0.15684631 |
| hsa04659: Th17 cell differentiation | 0.00734264 | 0.04807682 |
| hsa04658: Th1 and Th2 cell differentiation | 0.01032726 | 0.06173908 |
| hsa05323: Rheumatoid arthritis | 0.01092655 | 0.06393195 |
| GO: 0030669~clathrin-coated endocytic vesicle membrane | 0.01841431 | 0.42813264 |
| GO: 0010008~endosome membrane | 0.05188268 | 0.86361666 |
| GO: 0031901~early endosome membrane | 0.05200272 | 0.86361666 |
| GO: 0055038~recycling endosome membrane | 0.07280561 | 1 |
| GO: 0032588~trans-Golgi network membrane | 0.07280561 | 1 |
| GO: 0050852~T cell receptor signaling pathway | 0.24866213 | 1 |
| hsa05203: Viral carcinogenesis | 0.28823311 | 1 |
| hsa04218: Cellular senescence | 0.38971633 | 1 |
| hsa04144: Endocytosis | 0.49025631 | 1 |
| Annotation Cluster 4: Enrichment Score: 2.5719744078830575 |  |  |
| Term | PValue | Benjamini |
| GO: 0016339~calcium-dependent cell-cell adhesion via plasma membrane cell adhesion molecules | 4.69E-06 | 5.39E-04 |
| GO: 0007416~synapse assembly | 1.79E-04 | 0.015208 |
| GO: 0007268~chemical synaptic transmission | 0.12291758 | 1 |
| GO: 0045202~synapse | 0.49932314 | 1 |
| Annotation Cluster 5: Enrichment Score: 1.3966018993019305 |  |  |
| Term | PValue | Benjamini |
| hsa00480: Glutathione metabolism | 0.00361814 | 0.02618394 |
| GO: 0004364~glutathione transferase activity | 0.00366572 | 0.29096645 |
| hsa05204: Chemical carcinogenesis - DNA adducts | 0.00922531 | 0.05637689 |
| GO: 0006749~glutathione metabolic process | 0.02149396 | 1 |
| hsa00982: Drug metabolism - cytochrome P450 | 0.04022589 | 0.22124239 |
| hsa01524: Platinum drug resistance | 0.04229977 | 0.22808699 |
| hsa05207: Chemical carcinogenesis - receptor activation | 0.05243462 | 0.27729846 |
| hsa00980: Metabolism of xenobiotics by cytochrome P450 | 0.0536251 | 0.27824345 |
| hsa00983: Drug metabolism - other enzymes | 0.05860336 | 0.29301679 |
| GO: 0006805~xenobiotic metabolic process | 0.07181652 | 1 |
| GO: 0042178~xenobiotic catabolic process | 0.09474298 | 1 |
| hsa05225: Hepatocellular carcinoma | 0.15260351 | 0.68796663 |
| hsa05418: Fluid shear stress and atherosclerosis | 0.15821423 | 0.69977247 |
| hsa05208: Chemical carcinogenesis - reactive oxygen species | 0.23136469 | 0.92210563 |
| Annotation Cluster 6: Enrichment Score: 1.3570585896445353 |  |  |
| Term | PValue | Benjamini |
| GO: 0045869~negative regulation of single stranded viral RNA replication via double stranded DNA intermediate | 0.00518358 | 0.33745123 |
| GO: 0010529~negative regulation of transposition | 0.01639338 | 0.89908234 |
| GO: 0070383~DNA cytosine deamination | 0.02040899 | 0.99646883 |
| GO: 0047844~deoxycytidine deaminase activity | 0.02262817 | 1 |
| GO: 0009972~cytidine deamination | 0.02905456 | 1 |
| GO: 0016554~cytidine to uridine editing | 0.02905456 | 1 |
| GO: 0004126~cytidine deaminase activity | 0.03216004 | 1 |
| GO: 0080111~DNA demethylation | 0.08096562 | 1 |
| hsa03250: Viral life cycle - HIV-1 | 0.49555146 | 1 |
| GO: 0000932~P-body | 0.62879617 | 1 |
| Annotation Cluster 7: Enrichment Score: 1.2134687565396018 |  |  |
| Term | PValue | Benjamini |
| GO: 0045324~late endosome to vacuole transport | 0.01657295 | 0.89908234 |
| GO: 0097352~autophagosome maturation | 0.06918992 | 1 |
| GO: 0016236~macroautophagy | 0.1995957 | 1 |

**Table S3.**
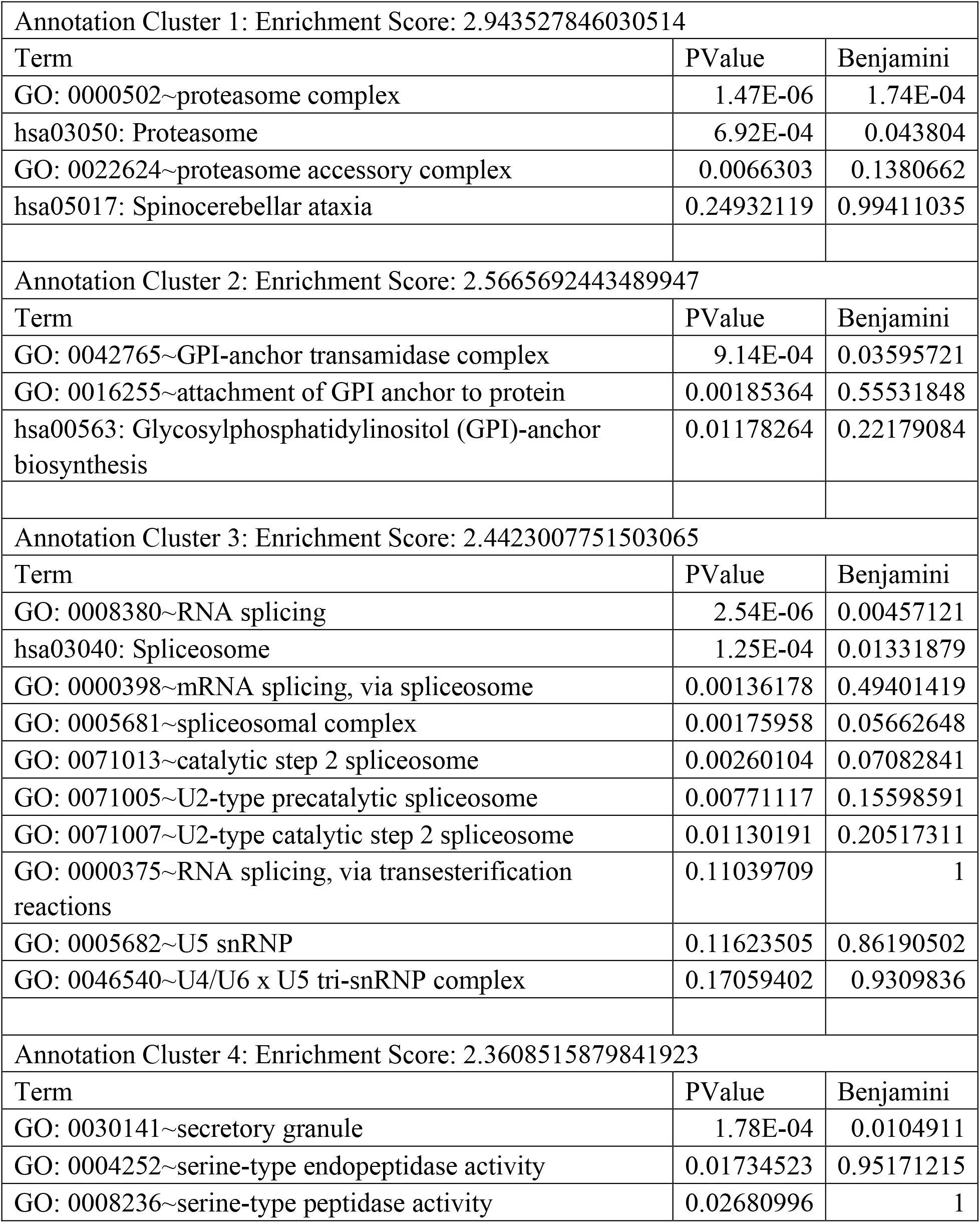
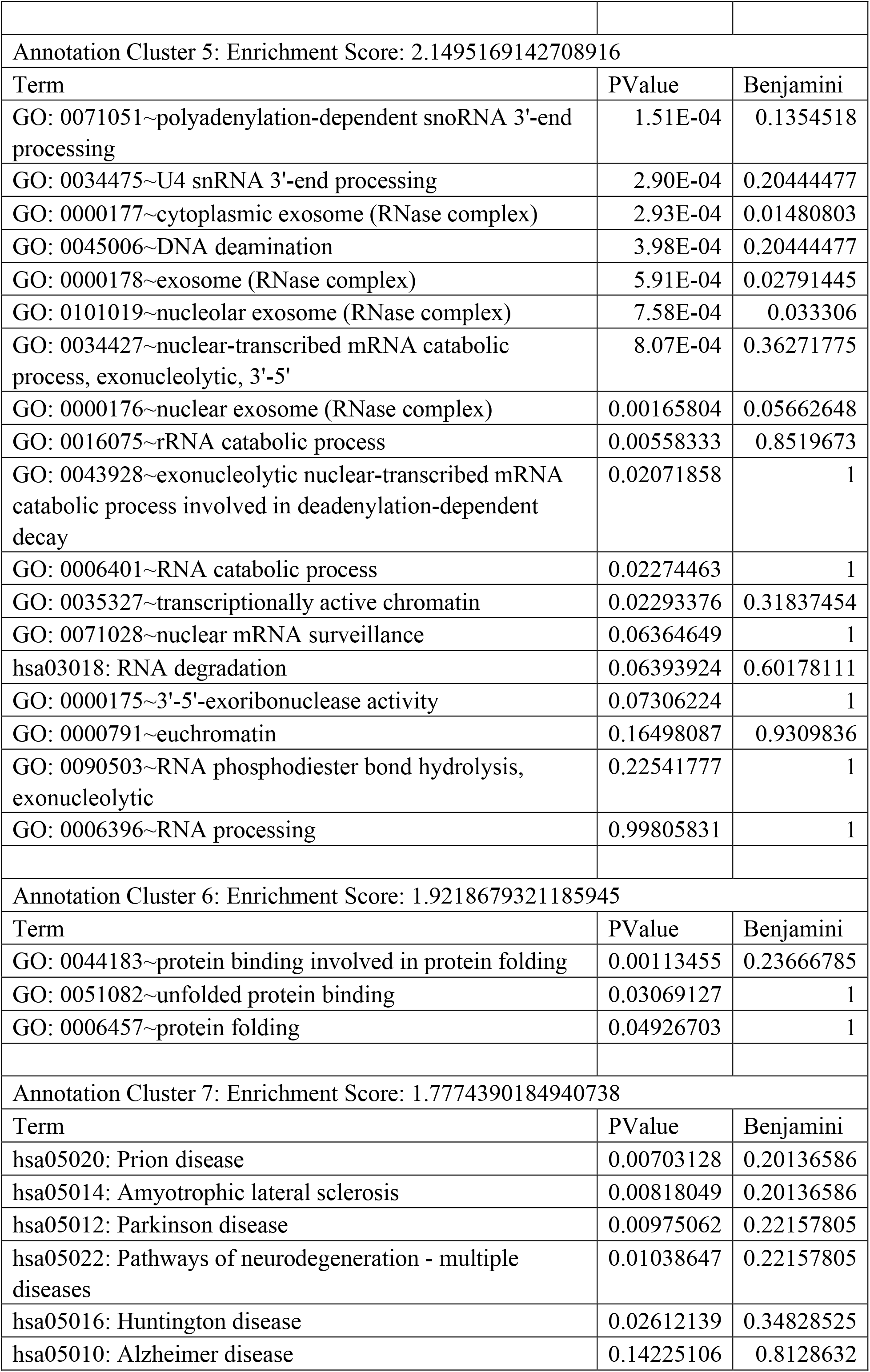
Functional annotation clustering of HCT116 structural-related intra-chromosomal PPIs.

|  |  |  |
| --- | --- | --- |
| Annotation Cluster 1: Enrichment Score: 2.943527846030514 |  |  |
| Term | PValue | Benjamini |
| GO: 0000502~proteasome complex | 1.47E-06 | 1.74E-04 |
| hsa03050: Proteasome | 6.92E-04 | 0.043804 |
| GO: 0022624~proteasome accessory complex | 0.0066303 | 0.1380662 |
| hsa05017: Spinocerebellar ataxia | 0.24932119 | 0.99411035 |
| Annotation Cluster 2: Enrichment Score: 2.5665692443489947 |  |  |
| Term | PValue | Benjamini |
| GO: 0042765~GPI-anchor transamidase complex | 9.14E-04 | 0.03595721 |
| GO: 0016255~attachment of GPI anchor to protein | 0.00185364 | 0.55531848 |
| hsa00563: Glycosylphosphatidylinositol (GPI)-anchor biosynthesis | 0.01178264 | 0.22179084 |
| Annotation Cluster 3: Enrichment Score: 2.4423007751503065 |  |  |
| Term | PValue | Benjamini |
| GO: 0008380~RNA splicing | 2.54E-06 | 0.00457121 |
| hsa03040: Spliceosome | 1.25E-04 | 0.01331879 |
| GO: 0000398~mRNA splicing, via spliceosome | 0.00136178 | 0.49401419 |
| GO: 0005681~spliceosomal complex | 0.00175958 | 0.05662648 |
| GO: 0071013~catalytic step 2 spliceosome | 0.00260104 | 0.07082841 |
| GO: 0071005~U2-type precatalytic spliceosome | 0.00771117 | 0.15598591 |
| GO: 0071007~U2-type catalytic step 2 spliceosome | 0.01130191 | 0.20517311 |
| GO: 0000375~RNA splicing, via transesterification reactions | 0.11039709 | 1 |
| GO: 0005682~U5 snRNP | 0.11623505 | 0.86190502 |
| GO: 0046540~U4/U6 x U5 tri-snRNP complex | 0.17059402 | 0.9309836 |
| Annotation Cluster 4: Enrichment Score: 2.3608515879841923 |  |  |
| Term | PValue | Benjamini |
| GO: 0030141~secretory granule | 1.78E-04 | 0.0104911 |
| GO: 0004252~serine-type endopeptidase activity | 0.01734523 | 0.95171215 |
| GO: 0008236~serine-type peptidase activity | 0.02680996 | 1 |
| Annotation Cluster 5: Enrichment Score: 2.1495169142708916 |  |  |
| Term | PValue | Benjamini |
| GO: 0071051~polyadenylation-dependent snoRNA 3'-end processing | 1.51E-04 | 0.1354518 |
| GO: 0034475~U4 snRNA 3'-end processing | 2.90E-04 | 0.20444477 |
| GO: 0000177~cytoplasmic exosome (RNase complex) | 2.93E-04 | 0.01480803 |
| GO: 0045006~DNA deamination | 3.98E-04 | 0.20444477 |
| GO: 0000178~exosome (RNase complex) | 5.91E-04 | 0.02791445 |
| GO: 0101019~nucleolar exosome (RNase complex) | 7.58E-04 | 0.033306 |
| GO: 0034427~nuclear-transcribed mRNA catabolic process, exonucleolytic, 3'-5' | 8.07E-04 | 0.36271775 |
| GO: 0000176~nuclear exosome (RNase complex) | 0.00165804 | 0.05662648 |
| GO: 0016075~rRNA catabolic process | 0.00558333 | 0.8519673 |
| GO: 0043928~exonucleolytic nuclear-transcribed mRNA catabolic process involved in deadenylation-dependent decay | 0.02071858 | 1 |
| GO: 0006401~RNA catabolic process | 0.02274463 | 1 |
| GO: 0035327~transcriptionally active chromatin | 0.02293376 | 0.31837454 |
| GO: 0071028~nuclear mRNA surveillance | 0.06364649 | 1 |
| hsa03018: RNA degradation | 0.06393924 | 0.60178111 |
| GO: 0000175~3'-5'-exoribonuclease activity | 0.07306224 | 1 |
| GO: 0000791~euchromatin | 0.16498087 | 0.9309836 |
| GO: 0090503~RNA phosphodiester bond hydrolysis, exonucleolytic | 0.22541777 | 1 |
| GO: 0006396~RNA processing | 0.99805831 | 1 |
| Annotation Cluster 6: Enrichment Score: 1.9218679321185945 |  |  |
| Term | PValue | Benjamini |
| GO: 0044183~protein binding involved in protein folding | 0.00113455 | 0.23666785 |
| GO: 0051082~unfolded protein binding | 0.03069127 | 1 |
| GO: 0006457~protein folding | 0.04926703 | 1 |
| Annotation Cluster 7: Enrichment Score: 1.7774390184940738 |  |  |
| Term | PValue | Benjamini |
| hsa05020: Prion disease | 0.00703128 | 0.20136586 |
| hsa05014: Amyotrophic lateral sclerosis | 0.00818049 | 0.20136586 |
| hsa05012: Parkinson disease | 0.00975062 | 0.22157805 |
| hsa05022: Pathways of neurodegeneration - multiple diseases | 0.01038647 | 0.22157805 |
| hsa05016: Huntington disease | 0.02612139 | 0.34828525 |
| hsa05010: Alzheimer disease | 0.14225106 | 0.8128632 |

